# Unpredictable circadian rhythm disruption in Wistar rats: biological and behavioural changes reflecting bipolar disorder pathophysiology

**DOI:** 10.1101/2025.09.06.674601

**Authors:** Heather K. Macpherson, Roger B. Varela, Sebastian C. McCullough, Tristan G. Houghton, Isha Chawla, Ning Wang, Venea D. Daygon, Xiaoying Cui, Susannah J. Tye

## Abstract

Circadian disturbances are implicated in dysregulation of arousal and general neurobiological function, contributing to conditions such as bipolar disorder (BD). However, the behavioural and biological consequences of circadian disruption on arousal dysfunction remain poorly quantified. Here, we developed a novel unpredictable circadian disruption (UCD) protocol – consisting of unpredictable exposure to light and sound – to investigate its impact on locomotor activity and its association with metabolic, inflammatory, stress, and circadian markers in the nucleus accumbens (NAc). Forty-eight Wistar rats were exposed to UCD or control conditions, with or without corticosterone administration, for five weeks. Body weight was tracked throughout the study. Locomotor activity was assessed over the final two weeks (nine sessions) in an open-field arena. Real-time PCR was used to quantify NAc gene expression of inflammatory, metabolic, stress, and circadian markers, while liquid chromatography-mass spectrometry (LC-MS) measured neurotransmitter and central carbon metabolite concentrations. UCD animals exhibited initial hyperactivity at three weeks, followed by hypoactivity at four weeks. UCD was associated with increased NAc expression of inflammatory, stress, and circadian markers. Male UCD animals showed significant weight gain, an effect reversed in females. UCD also induced increases in NAc insulin resistance markers and reductions in central carbon metabolites, indicating disrupted striatal glucose metabolism. These findings highlight the central effects of circadian disruption on locomotor behaviour, stress, and immunometabolic signalling, offering mechanistic insights into arousal dysfunction in BD.

**Highlights:**

- Unpredictable circadian disruption (UCD) triggers a biphasic shift in locomotor behaviour
- UCD alters levels of striatal inflammatory, circadian, stress, and metabolic markers
- UCD has sex-specific effects on behaviour, weight, and molecular biomarkers
- Corticosterone administration mitigates key molecular disruptions induced by UCD

## 1 Introduction

Bipolar disorder (BD) is a serious psychiatric condition characterised by mood disturbances, including episodes of depression presenting as low mood, fatigue, psychomotor retardation, and reduced energy; and mania, which can include heightened euphoria or dysphoria, agitation, impulsivity, and hyperactivity (McIntyre et al., 2020). Alongside severe mood dysregulation and other psychiatric symptoms such as psychosis, BD is associated with physical changes, such as circadian rhythm dysregulation, inflammation, hypothalamic-pituitary-adrenal (HPA) axis hyperactivity, impaired glucose metabolism, and disrupted energy homeostasis (McIntyre et al., 2020; Rosenblat et al., 2015). Pharmacological treatments remain the cornerstone of BD management; however, current medications often provide only partial symptom relief, even when following best-practice guidelines (Novick and Swartz, 2019). A major obstacle to improved therapeutics is the limited understanding of BD pathophysiology. Developing treatments that target the underlying causal mechanisms of mood disorders is critical for achieving complete symptom resolution and preventing relapse.

Several systemic disturbances have been implicated in BD pathophysiology, including immune dysregulation, mitochondrial dysfunction, and impaired insulin signalling (McIntyre et al., 2020). Another key contributor is circadian disturbance, which manifests as poor sleep quality, increased sleep variability, prolonged sleep latency, daytime dysfunction, and increased wakefulness after sleep onset. These abnormalities are consistently observed across mood states, demonstrating that circadian dysfunction is a core trait of BD (Bradley et al., 2017; Cretu et al., 2016; Harvey et al., 2005; Krishnamurthy et al., 2018; Mehl et al., 2006; Menkes et al., 2021; Millar et al., 2004; Roloff et al., 2022). This circadian disturbance has been observed in euthymic, manic, depressed and mixed mood states, thus making it a trait phenotype of BD (Bradley et al., 2017; Cretu et al., 2016; Giglio et al., 2010; Godin et al., 2017; Harvey et al., 2005; Krishnamurthy et al., 2018; Laskemoen et al., 2019; Menkes et al., 2021; Millar et al., 2004; Saunders et al., 2015). Circadian-disturbed BD patients frequently demonstrate more severe symptoms, a greater risk of recurrence, worsened functional impairment, longer duration of illness, and a lessened likelihood of successful treatment (Cretu et al., 2016; Giglio et al., 2010; Godin et al., 2017; Lai et al., 2014; Laskemoen et al., 2019; Lunsford-Avery et al., 2012; Menkes et al., 2021; Saunders et al., 2015; Sylvia et al., 2018; Sylvia et al., 2012). Additionally, high-risk patients with circadian disturbance have been shown to have a much higher probability of developing BD than high-risk patients with normal circadian rhythm (Levenson et al., 2017; Ritter et al., 2015; Soehner et al., 2019).

The mechanisms linking circadian disruption to BD are thought to involve dysregulated limbic system transmission mediated by clock genes, HPA axis dysfunction, and altered melatonergic signalling (Ketchesin et al., 2020). Circadian dysfunction has also been implicated in BD-associated metabolic comorbidities due to its essential role in regulating glucose homeostasis, insulin sensitivity, and energy balance (Yan et al., 2023). Moreover, it has been proposed that disrupted sleep patterns drive neuroinflammation through impaired circadian regulation of immune responses and altered metabolic clearance in the brain (Dallaspezia and Benedetti, 2015). Clinical studies further link sleep disturbances to type 2 diabetes and obesity, highlighting the broad physiological consequences of circadian disruption (Miller and Howarth, 2023).

Due to its complex phenotype and unknown pathophysiology, BD is an exceedingly difficult condition to research, which reflects the lack of effective, translational animal models of the condition (Beyer and Freund, 2017). A key feature of BD that is rarely recapitulated in animal models, for instance, is a “mood-switch” (i.e., a transition from mania-like to depressive-like behaviours, or vice versa) (Beyer and Freund, 2017). A beneficial method of investigating BD lies within the Research Domain Criteria (RDoC) framework, which views psychopathology as deviations from normal neurobiological functions, and aims to understand how disturbances of these functions can contribute to psychiatric symptomatology (Anderzhanova et al., 2017). The neurobiological domains defined by RDoC – cognitive systems, negative valence systems, positive valence systems, social processes, arousal/regulation, and sensorimotor systems – are well conserved through mammalian evolution, which strengthens translational validity between animal and clinical studies that utilise this framework (Anderzhanova et al., 2017; Sanislow et al., 2019). RDoC emphasises the usage of both behavioural (e.g., locomotor activity) and biological (e.g., metabolomics) assessments in order to comprehensively detail mechanisms underpinning neurobiological functioning along the continuum from normal to pathological (Sanislow et al., 2019).

Of particular relevance to BD is the RDoC domain of arousal/regulation, which includes systems of arousal, circadian rhythm, and sleep-wakefulness (Anderzhanova et al., 2017). A key feature of arousal is modulation in activity and energy levels; thus, dysregulated arousal can result in the transdiagnostic behavioural endophenotypes of hyperactivity, or hypoactivity (NIMH, 2016). Given the relationship between circadian disturbance and mood dysregulation in BD, as well as the potential involvement of other biological mechanisms underlying this phenotype, the present study aimed to examine the effects of a novel unpredictable circadian disruption (UCD) protocol on the arousal system. Specifically, we examined UCD-induced changes in locomotor activity, metabolic and inflammatory markers, stress responses, and circadian rhythm alterations within the limbic system. Additionally, we explored sex-dependent effects of UCD on these variables and assessed its interaction with HPA axis activation to further elucidate the biological mechanisms underlying arousal dysregulation in BD.

## 2 Methods

### 2.1 Ethics

All animal protocols were approved by the University of Queensland Animal Ethics Committee (Protocol number: 2022/AE000132). All experimental procedures were performed in accordance with the NIH and NHMRC codes for the care and use of animals for scientific purposes, and all efforts were made to reduce the number of animals used.

### 2.2 Animals

Forty-eight albino Wistar rats (24 male and 24 female), 45 days old weighing 155-225g (female) and 198-259g (male) at the beginning of the study were provided by the University of Queensland Biological Resources animal facility. Animals were single-housed, at a temperature of 22±1°C, with food and water available *ad libitum*. Animal health and weight were measured regularly.

### 2.3 Experimental protocol

#### Chronic unpredictable sleep disruption

A novel protocol designed by the authors was administered over five weeks for 24 animals. Non-UCD animals (n=24) were maintained under a twelve-hour light/dark cycle (lights on at 0600 hours). UCD animals were exposed to eight conditions designed to disturb sleep throughout both the light (0600 to 1800) and dark (1800 to 0600) periods. One condition for the light period and one for the dark period were selected randomly each day to ensure unpredictability (see Table 1 below, and Supplementary Data 1 for more details).

**Table 1:** Description of unpredictable circadian disruption (UCD) conditions.

| **Light period conditions** | **Dark period conditions** |
| --- | --- |
| Lights turned on/off every two hours. | Strobe light set at 120 flashes per minute. |
| Lights turned on/off every three hours. | Lights turned on/off every three hours. |
| Light period shortened to six hours. | Dark period shortened to six hours. |
| 65-80dB noise (traffic, construction, and airport noise) for twelve hours. | 65-80dB noise (traffic, construction, and airport noise) for twelve hours. |

#### Administration of corticosterone via drinking water

Rats were exposed to vehicle (n=24, twelve females) or corticosterone (n=24, twelve females) for 35 days. Corticosterone powder (Merck; Cat:#C2505-500MG) was dissolved in 100% ethanol and then added to drinking water to a final concentration of 200μg/mL and 1% ethanol. Vehicle was 1% ethanol in drinking water. Water was supplemented with freshly prepared vehicle or corticosterone every few days. Corticosterone bottles were covered in foil to protect against light-induced degradation. Thus, the final experiment consisted of eight groups, each with six animals: Corticosterone/UCD/Female, Corticosterone/Non-UCD/Female, Corticosterone/UCD/Male, Corticosterone/Non-UCD/Male, Vehicle/UCD/Female, Vehicle/non-UCD/Female, Vehicle/UCD/Male, and Vehicle/Non-UCD/Male.

### 2.4 Behavioural testing

Behavioural testing began after 21 days of UCD or sham (non-UCD) treatment. Open field test (OFT) trials were repeated nine times, once every day for five days in the first week and four days in the second. Testing was conducted between 9 and 11am in black 60×60×60 cm arenas under diffuse bright light, in a quiet room. After the animal was placed in the top right corner of the arena, locomotor activity over a five-minute period was recorded on a top-mounted camera. Ethovision XT 16 (Noldus) was used to measure time in centre, latency to centre, centre-crossing frequency, and distance travelled. Rearing and grooming behaviours were manually scored. The tenth behavioural session was excluded from analysis.

### 2.5 Dissection and euthanasia

Rats were euthanised immediately after the final OFT session with intraperitoneal injections of 20μg Lethabarb (Virbac Animal Health; Cat:#1PO643-1). Blood was collected via cardiac puncture into 15ml tubes containing 100 units of heparin, and samples were centrifuged at 1200xg for 10 minutes. Plasma was removed from the top aqueous layer and stored at −80°C for posterior analysis. Brains were collected, snap-frozen in liquid nitrogen and stored at −80°C. Frozen brains were sectioned (1mm sections) and bilateral punch samples of nucleus accumbens (NAc) – the central component of the limbic system – were collected according to rat brain atlas coordinates (total of eight 1mm punches per animal) (Paxinos and Watson, 2014). Four punches each were allocated to gene expression analysis or metabolomics / neurotransmitter quantification.

### 2.6 Nucleus accumbens gene expression analysis

RNA was extracted from NAc brain punches using the RNeasy^®^ Mini kit (QIAGEN; Cat:#74106) and the RNAse-free DNAse kit (QIAGEN; Cat:#79256) according to the manufacturer’s instructions. Two samples were excluded from further analyses due to low RNA concentrations. 400ng total RNA per sample was used for cDNA synthesis. cDNA was synthesised from total RNA using the SensiFAST™ cDNA synthesis kit (Bioline; Cat:#BIO- 65054), and RT-PCR was performed using the SensiFAST™ SYBR Green Master Mix (Bioline; Cat:#BIO-98020) according to the manufacturer’s instructions. The following primers were used: HPRT (used as endogenous control), PRKAA2, MAPK1, TNFA, IL1B, NFKB1, INSR, PEPCK, TH, NR3C1, MTNR1A, MTNR1B, and CLOCK (see Supplementary Table 1). Each sample was tested in duplicate. RT-PCR consisted of a denaturation step (95°C for 10 minutes), then amplification (95°C for 15 s, 60°C for 20s and 72°C for 30s) for 40 cycles, performed in the LightCycler^®^ 480 Instrument II (Roche Diagnostics; Cat:#05015278001). The ΔCt method was used to assess gene expression.

### 2.7 Nucleus accumbens neurotransmitter and central carbon metabolite (CCM) analysis

#### 2.7.1 Extraction of metabolites and neurotransmitters

1ml 50% ice cold methanol and 0.50µM internal standard were added to NAc tissue punch samples and sonicated for ten minutes in ice water. 300µl chloroform was added and the samples were centrifuged at 16000xg for five minutes at 4°C. The top aqueous layer was removed and the methanol evaporated using a vacuum concentrator. The leftover water fraction was freeze dried overnight, reconstituted in 100µl 2% acetonitrile solution, then centrifuged at 16000xg for five minutes at 4°C. All standards were serially diluted from 200µM to 1.5nM and added with 5µM azidothymidine.

#### 2.7.2 Central carbon metabolite (CCM) analysis

CCM (see Supplementary Table 2) levels were quantified from samples extracted from NAc tissue using liquid chromatography-tandem mass spectrometry (LC-MS/MS). LC-MS/MS was performed using a Nexera ultra high-performance liquid chromatograph (Shimadzu) coupled to the 8060 triple quadrupole mass spectrometer (Shimadzu) on Acquity HSS T3 1.8μm, 2.1×100mm columns (Waters Corporation; Cat:#186003539) with Acquity HSS T3 1.8µm 2.1×5mm guard columns (Waters Corporation; Cat:#186003976). LC-MS/MS was performed on negative ionisation mode with a 250µl/min flow rate, 10µl injection volume, and column temperature of 45°C. 7.5mM tributylamine with pH adjusted to 4.95 with acetic acid was used for mobile phase A, and 100% acetonitrile for mobile phase B. Total analysis time per sample was thirty-five minutes.

#### 2.7.3 Neurotransmitter quantification

Dopamine, serotonin, acetylcholine, and gamma-aminobutyric acid (GABA) levels were quantified from samples extracted from NAc tissue using liquid chromatography-mass spectrometry (LC-MS). Melatonin was not quantifiable due to low concentration. LC-MS was performed using the same instrument described above on Gemini 3µm NX-C18 15×2mm columns (Phenomenex; Cat:#00F-4453-B0) with Gemini NX 2×4mm guard columns (Phenomenex; Cat:#AJ0-8367). LC-MS was performed with a 300µl/min flow rate, 10µl injection volume, and column temperature of 40°C. 0.1% formic acid in water was used for mobile phase A and 0.1% formic acid in acetonitrile for mobile phase B. Total analysis time per sample was sixteen minutes. Multiple reaction monitoring (MRM) transitions used to quantify the neurotransmitters are given in Supplementary Table 3.

### 2.9 Molecular assays using plasma

#### 2.9.1 Protein concentration

Plasma sample protein concentration was measured by Bradford Protein Assay (Biorad; Cat:#5000204), according to the manufacturer’s instructions. Each aliquot used were measured independently. Values were used to standardise ELISA results and glucose levels.

#### 2.9.2 Enzyme-linked immunosorbent assays (ELISA)

Plasma levels of insulin and corticosterone were determined using commercially available ELISA kits (Invitrogen; Cat:#ERINS and #EIACORT) according to the manufacturer’s instructions. Samples were diluted eight-fold (for the insulin ELISA) and 100-fold (for corticosterone) in the provided assay diluents. All samples and standards were run in duplicate. Absorbance was measured using the CLARIOstar Plus microplate reader (BMG LABTECH GmbH).

#### 2.9.3 Measurement of glucose levels

20µl of plasma was applied to a test strip and inserted into the Accu-Chek Instant S blood glucose meter (Roche Diabetes Care). Samples were tested in duplicate.

### 2.10 Statistical analysis

All statistical analyses were performed on GraphPad Prism 10.0.2 (GraphPad Software Inc.), except for multivariate analysis, which was performed with SIMCA 18.0.1 (Umetrics, Sartorius). Normality of data was assessed using the Shapiro-Wilk and D’Agostino-Pearson tests, and heteroscedasticity with Spearman’s test. Normal data were analysed using two-way analysis of variance (ANOVA) or three-way ANOVA with Geisser-Greenhouse correction, and on non-normal data using multiple Mann-Whitney tests followed by Holm-Šidak correction. Post-hoc tests were not performed on ANOVA analyses due to low power. Longitudinal analyses of behavioural data were conducted using Spearman’s correlation. Simple linear regression analyses with Holm-Šídák correction were used to examine relationships between biological and behavioural variables. Multivariate analysis was performed using orthogonal partial-least squares discriminant analysis (OPLS-DA). Further details of statistical analyses can be found in Supplementary Data 2.

## 3 Results

### 3.1 Effects of unpredictable circadian disruption on locomotor behaviour

A significant effect of UCD was found on time in centre [F(1,38)=7.491, p=0.0094], centre-crossing frequency [F(1,38)=5.865, p=0.0203], and rearing [F(1,38)=4.139, p=0.0489] on the first day of testing, with UCD-treated animals showing increases in these parameters when compared to non-UCD animals (see Figure 1A-C). An effect of corticosterone was also found on distance travelled [F(1,38)=4.350, p=0.0438] and rearing [F(1,38)=4.954, p=0.0320] on day one of testing (see Figure 1C-D). Subsequently, UCD animals exhibited significant decreases in rearing on days six [F(1,39)=7.182, p=0.0107], seven [F(1,36)=17.08, p=0.0002], eight [F(1,40)=9.585, p=0.0036], and nine (final) [F(1,40)=21.58, p<0.0001] of testing when compared to non-UCD controls; as well as distance travelled [F(1,40)=6.474, p=0.0149] on the final day of testing (see Figure 1E-F and Supplementary Figure 1A-C). When behaviours were analysed longitudinally, it was found that UCD animals had significant negative linear habituation patterns for distance travelled [UCD/Corticosterone/Female: r=-0.850, p=0.0061; UCD/Corticosterone/Male: r=-0.900, p=0.0020; UCD/Vehicle/Female: r=-0.900, p=0.0020; UCD/Vehicle/Male: r=-0.867, p=0.0045] and rearing [UCD/Corticosterone/Female: r=-0.900, p=0.0020; UCD/Corticosterone/Male: r=-0.833, p=0.0083; UCD/Vehicle/Female: r=-0.933, p=0.0007; UCD/Vehicle/Male: r=-0.717, p=0.0369] that were not seen in non-UCD controls (see Figure 1G-J). Overall, these results suggest that UCD animals expressed a hyperactive behavioural phenotype after three weeks of treatment, followed by exacerbated behavioural habituation beginning at four weeks, and progressing to a hypoactive phenotype in the last session.

**Figure 1:**
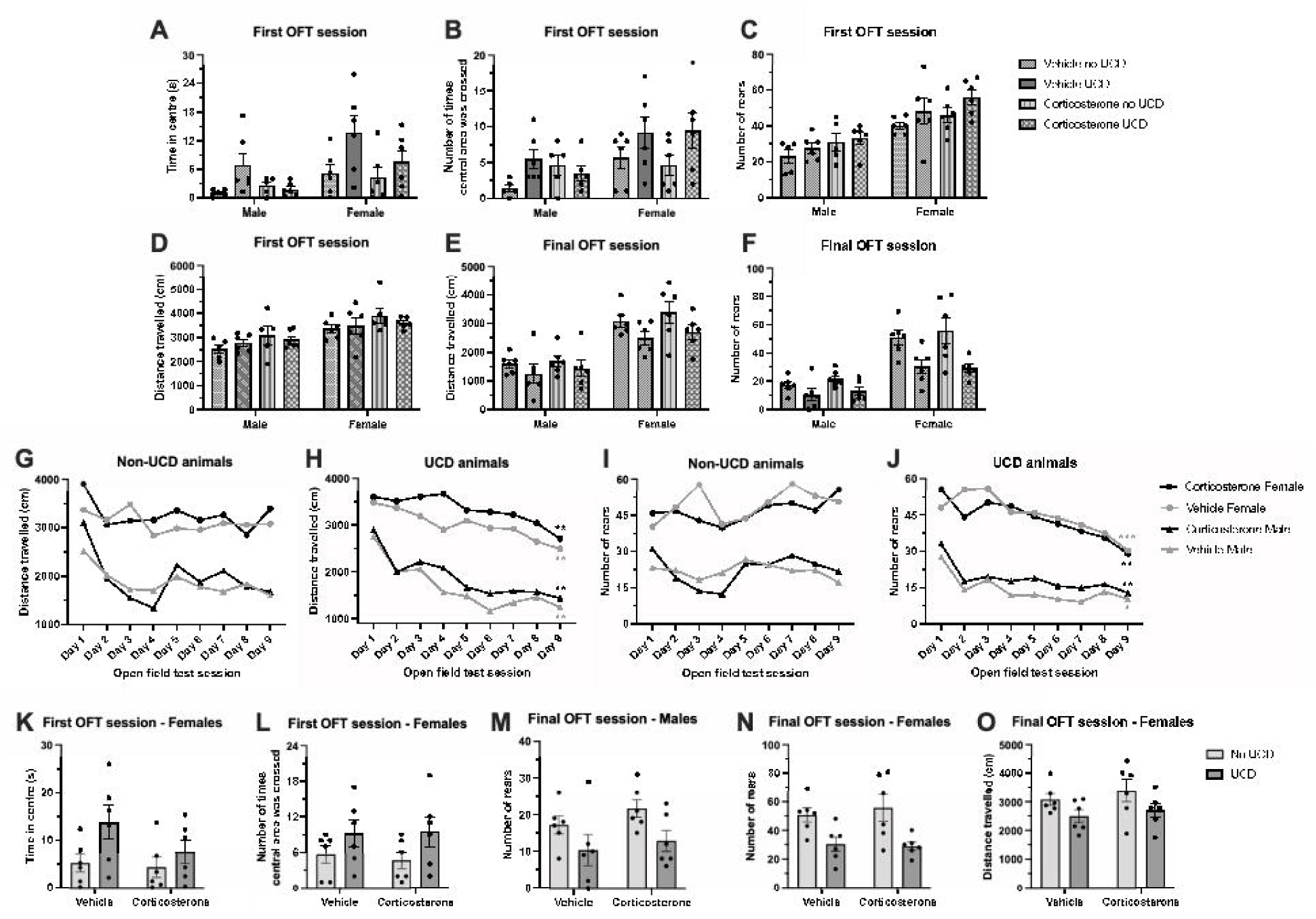
Effects of unpredictable circadian disturbance (UCD) on locomotor behaviour. Mean (±SEM) time in centre (seconds) ***(A)***, centre-crossing frequency ***(B)***, number of rears ***(C)***, and distance travelled (cm) ***(D)*** on the first day of testing; and distance travelled ***(E)*** and number of rears ***(F)*** on the final day of testing, compared between UCD/non-UCD, corticosterone/vehicle, and male/female groups. Effect of UCD is p < 0.05 for A-C and E-F; and effect of corticosterone is p < 0.05 for C-D, as measured by three-way analysis of variance (ANOVA) with Geisser-Greenhouse correction. Longitudinal models of distance travelled (cm) per session in non-UCD ***(G)*** and UCD ***(H)*** animals; and number of rears per session in non-UCD ***(I)*** and UCD ***(J)*** animals. *p < 0.05, **p < 0.01, ***p < 0.001 as measured using Spearman’s correlation of time (session) against locomotor behaviour (distance or rearing). Mean (±SEM) time in centre (seconds) ***(K)*** and centre-crossing frequency ***(L)*** by females only on the first day of testing; number of rears by males only ***(M)*** and females only ***(N)*** on the final day of testing; and distance travelled by females only on the final day of testing ***(O)***. Effect of UCD is p < 0.05, as measured by two-way ANOVA with Geisser-Greenhouse correction.

UCD-treated females demonstrated significant increases in time in centre [F(1,21)=5.171, p=0.0336] and centre-crossing frequency [F(1,21)=4.677, p=0.0422] on the first day of testing (see Figure 1K-L). UCD males demonstrated significant decreases in rearing on days five [F(1,21)=5.072, p=0.0351], six [F(1,21)=7.086, p=0.0146], seven [F(1,21)=12.60, p=0.0019], eight [F(1,21)=4.368, p=0.0490] and the final day [F(1,21)=6.864, p=0.0160] of testing when compared to non-UCD males, but UCD females demonstrated significant decreases in rearing only on days seven [F(1,17)=6.909, p=0.0176], eight [F(1,21)=5.720, p=0.0262], and the final day [F(1,21)=15.81, p=0.0007] when compared to non-UCD females (see Figure 1M-N and Supplementary Figure 1D-I). UCD females also exhibited a decrease in distance travelled [F(1, 21)=5.652, p=0.0270] on the final day of testing when compared to non-UCD females (see Figure 1O). Overall, these results suggest that females may be more susceptible to UCD-induced hyperactivity than males, and UCD-treated males may progress to a hypoactive phenotype earlier than UCD-treated females.

### 3.2 Effects of unpredictable circadian disruption on inflammatory markers

Three-way ANOVAs were not able to be performed due to the non-normal nature of gene expression data. Instead, male and females were combined, and data were analysed using multiple Mann-Whitney tests with Holm-Šídák correction. Vehicle-treated UCD animals demonstrated significant increases in IL1B [U=10.00, p=0.0004], TNFA [U=16.00, p=0.0049], and an almost significant increase in AMPK [U=33.50, p=0.0893] gene expression in the NAc when compared to vehicle-treated non-UCD controls (see Figure 2A-C). When females and males were analysed separately, a significant effect of sex was found. Significant increases in IL1B [U=0.000, p=0.0086], TNFA [U=0.000, p=0.0158], and NFKB [U=0.000, p=0.0315] were found in vehicle-treated UCD males when compared to vehicle-treated non-UCD males (see Figure 2D-F). These results suggest that inflammation in the NAc is significantly increased in UCD-treated animals when compared to controls, and that the effect is exacerbated in UCD-treated males.

**Figure 2:**
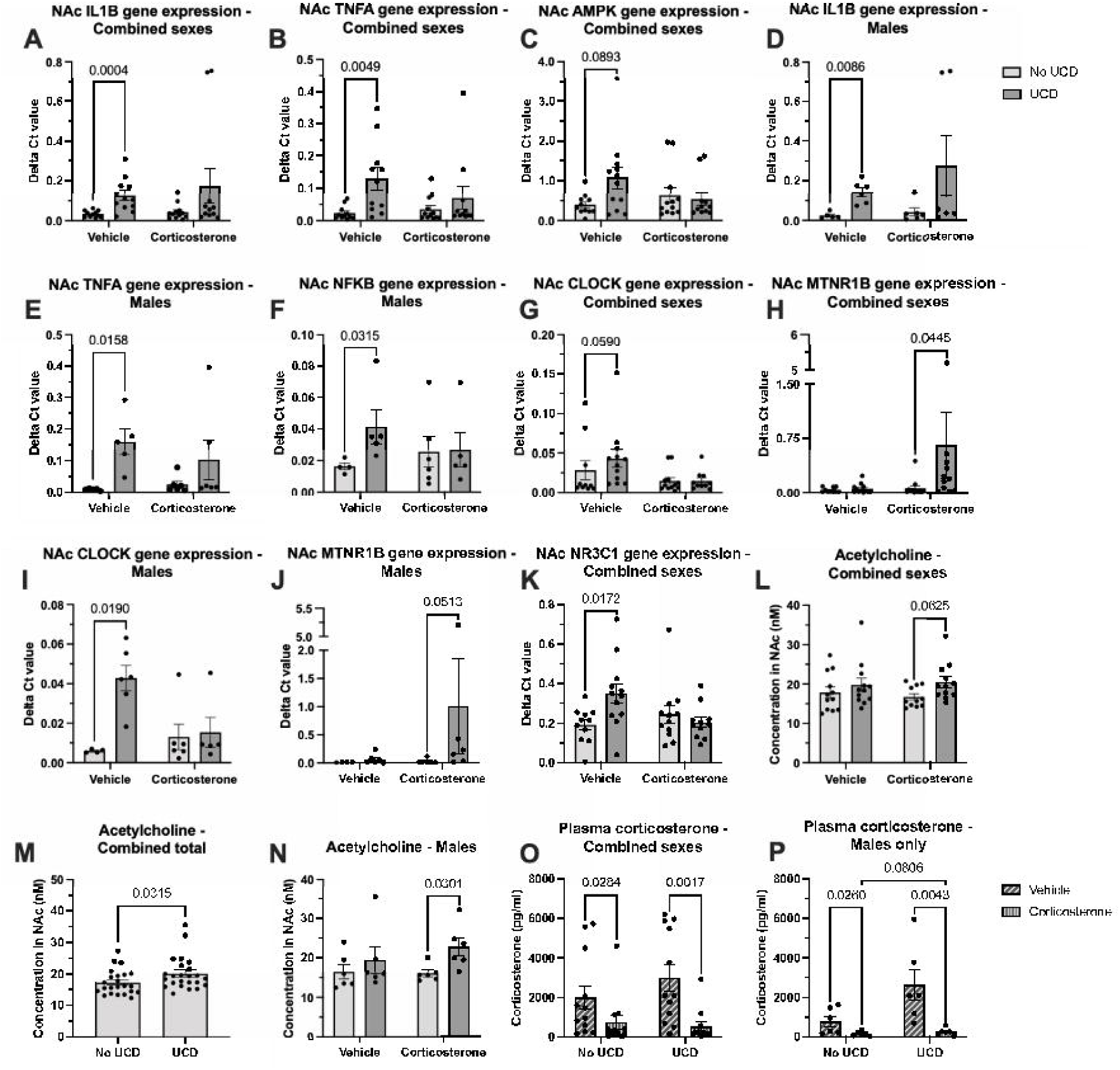
Effects of unpredictable circadian disturbance (UCD) on nucleus accumbens (NAc) neurotransmitter concentrations, and markers of inflammation, circadian rhythm, stress. Mean (±SEM) gene expression (expressed as ΔCt) in the NAc of IL1B *(A)* TNFA *(B)*, and AMPK *(C)* of males and females combined; IL1B *(D)*, TNFA *(E)*, and NFKB *(F)* in males only; CLOCK *(G)*, and MTNR1B *(H)* of males and females combined; CLOCK *(I)* and MTNR1B *(J)* of males only; and NR3C1 *(K)* of males and females combined. Mean (±SEM) NAc concentration of acetylcholine in males and females combined *(L)*; pooled UCD and non-UCD groups *(M)*; and males only *(N)*. Mean (±SEM) plasma concentration (pg/ml) of corticosterone in males and females combined *(O)* and males only *(P)*. Displayed p-values were calculated using multiple Mann-Whitney tests with Holm-Šídák correction.

### 3.3 Effects of unpredictable circadian disruption on circadian markers

Vehicle-treated UCD animals demonstrated an almost significant increase in NAc CLOCK [U=27.00, p=0.0590] gene expression in the NAc when compared to vehicle-treated non-UCD controls (see Figure 2G). Corticosterone-treated UCD animals demonstrated a significant increase in NAc MTNR1B gene expression [U=29.00, p=0.0445] when compared to corticosterone-treated non-UCD controls (see Figure 2H). A significant increase in NAc CLOCK [U=0.000, p=0.0190] expression was found in vehicle-treated UCD males when compared to vehicle-treated non-UCD males, and an almost significant increase in NAc MTNR1B [U=4.000, p=0.0513] expression was found in corticosterone-treated UCD males when compared to corticosterone non-UCD males (see Figure 2I-J).

### 3.4 Effects of unpredictable circadian disruption on stress markers

UCD treatment significantly increased NR3C1 gene expression in the NAc [U=24.00, p=0.0172], but only in vehicle-treated animals (see Figure 2K). UCD treatment had no effect on plasma corticosterone levels; however, multiple Mann-Whitney tests unveiled a significant increase in corticosterone in vehicle-treated animals when compared to corticosterone-treated animals, in both UCD [U=17.00, p=0.0017] and non-UCD [U=34.00, p=0.0284] animals (see Figure 2O). When sexes were analysed separately, vehicle-treated UCD [U=0.000, p=0.0043], and non-UCD [U=4.000, p=0.0260] males demonstrated significant increases in corticosterone levels when compared to corticosterone-treated males (see Figure 2P). An almost significant increase in plasma corticosterone was also seen in vehicle-treated UCD males [U=5.000, p=0.0806] when compared to vehicle-treated non-UCD males (see Figure 2P).

Multiple Mann-Whitney tests on data with sexes combined also unveiled an almost significant increase in NAc acetylcholine [U=31.00, p=0.0625] in corticosterone-treated UCD animals when compared to non-UCD corticosterone-treated controls (see Figure 2L). When the vehicle and corticosterone groups were pooled, a significant increase in NAc acetylcholine was found in UCD animals [U=175.0, p=0.0315] when compared to non-UCD animals (see Figure 2M). Corticosterone-treated UCD males exhibited significant increases in NAc acetylcholine [U=3.000, p=0.0301] when compared to corticosterone-treated non-UCD males (see Figure 2N).

### 3.5 Effects of unpredictable circadian disruption on metabolism

UCD males gained significantly more weight over the five-week period than non-UCD males [F(1,21)=11.49, p=0.0028], whereas UCD-treated females gained significantly less weight than their non-UCD counterparts [F(1,21)=5.866, p=0.0246] (see Figure 3A-B). Corticosterone was also found to have a significant suppressive effect on weight gain when all groups were compared [F(1,40)=65.03, p<0.0001] and in males only [F(1,21)=91.82, p<0.0001] (see Figure 3A and Supplementary Figure 2A).

**Figure 3:**
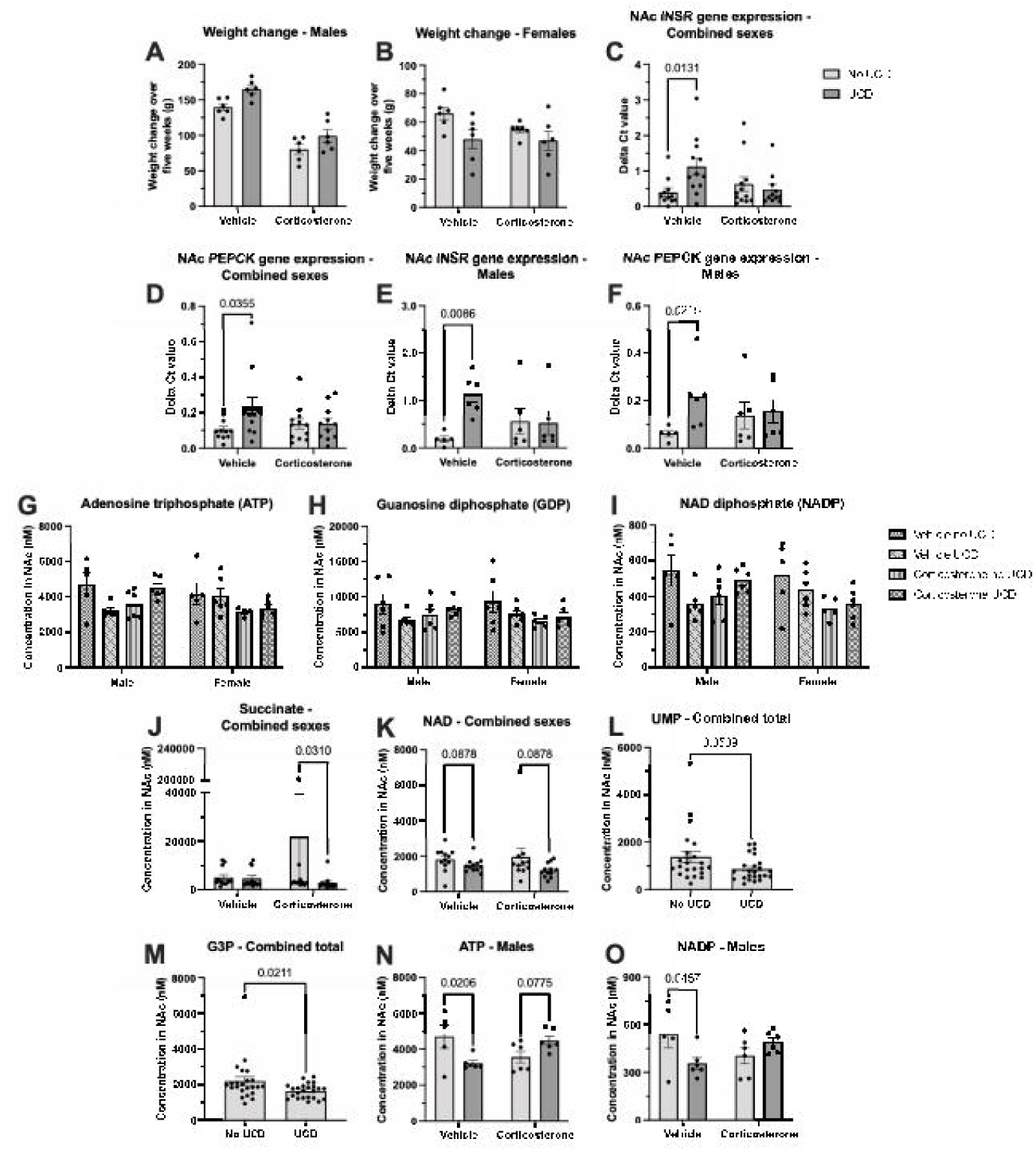
Effects of unpredictable circadian disturbance (UCD) on weight gain and markers of glucose metabolism in the nucleus accumbens (NAc). Mean (±SEM) weight change over five weeks (grams) of males only *(A)* and females only *(B)*, compared between UCD/non-UCD and corticosterone/vehicle groups. Effect of UCD is p < 0.05 for analyses, as measured by two-way ANOVA with Geisser-Greenhouse correction. Mean (±SEM) gene expression (expressed as ΔCt) in the NAc of INSR *(C)* and PEPCK *(D)* of males and females combined; and INSR *(E)* and PEPCK *(F)* of males only. Displayed p-values were calculated using multiple Mann-Whitney tests with Holm-Šídák correction. Mean (±SEM) NAc concentration (nM) of adenosine triphosphate (ATP) *(G)*, guanosine diphosphate (GDP) *(H)*, and nicotinamide adenine dinucleotide diphosphate (NADP) *(I)*, compared between UCD/non-UCD, corticosterone/vehicle, and male/female groups. Effect of UCD x Corticosterone is p < 0.05 for all analyses shown, as measured by three-way ANOVA with Geisser-Greenhouse correction. Mean (±SEM) NAc concentration (nM) of succinate *(J)*, oxidised nicotinamide adenine dinucleotide (NAD) *(K)*, and uridine monophosphate (UMP) *(L)* of males and females combined; and glycerol 3-phosphate (G3P) *(M)* of pooled UCD and non-UCD groups. Mean (±SEM) NAc concentration (nM) of ATP *(N)* and NADP *(O)* in males only. Displayed p-values were calculated using multiple Mann-Whitney tests with Holm-Šídák correction.

Vehicle-treated UCD animals demonstrated a significant increase in INSR [U=23.00, p=0.0131] and PEPCK [U=28.00, p=0.0355] gene expression in the NAc when compared to vehicle-treated non-UCD controls (see Figure 3C-D). Vehicle-treated UCD males also demonstrated increased NAc gene expression of INSR [U=0.000, p=0.0086] and PEPCK [U=1.500, p=0.0215] (see Figure 3E-F). No significant differences were found between groups in plasma insulin or glucose levels, neither when sexes were separated nor combined (see Supplementary Figure 2B-C).

Three-way ANOVAs unveiled a significant effect of Corticosterone x UCD on adenosine triphosphate (ATP) [F(1,36)=6.436, p=0.0157], guanosine diphosphate (GDP) [F(1,39)=4.858, p=0.0335], and nicotinamide adenine dinucleotide (NAD) phosphate [F(1,37)=6.515, p=0.0150] concentration in the NAc (see Figure 3G-I). Analysis of metabolomics data by multiple Mann-Whitney or t-tests with sexes combined further revealed significant reductions in GDP [t=2.342, difference between means (MD)±SEM=2004±855.8, p=0.0472] and NADP [t=2.470, MD±SEM=134.0±54.23, p=0.0352] in vehicle-treated UCD animals when compared to non-UCD vehicle controls, and succinate [U=27.00, p=0.0310] in corticosterone-treated UCD animals when compared to corticosterone non-UCD controls; and almost significant decreases in oxidised NAD in corticosterone-treated [U=37.00, p=0.0878], and vehicle-treated [U=37.00, p=0.0878] UCD animals when compared to their corresponding non-UCD controls (see Figure 3J-K and Supplementary Figure 2D-E). Analysis with pooled UCD and non-UCD groups revealed further significant decreases in oxidised NAD [U=156.0, p=0.0101] and glycerol-3-phosphate [U=168.0, p=0.0211], and an almost significant decrease in uridine monophosphate [U=184.0, p=0.0509] in UCD animals when compared to non-UCD controls (see Figure 3L-M and Supplementary Figure 2F). Results from metabolomics suggest an overall depressive effect of UCD on energy production in the NAc, with a potential modulating effect of corticosterone.

In males, two-way ANOVA identified a significant effect of UCD x Corticosterone on ATP [F(1,19)=11.21, p=0.0034] and NADP [F(1,19)=6.806, p=0.0173] concentration in the NAc (see Figure 3N-O). Further analysis of these data with multiple unpaired t-tests highlighted a significant decrease in ATP [t =2.846, MD±SEM=1489±523.2, p=0.0206] and NADP [t=2.471, MD±SEM=186.6±75.51, p=0.0457] in vehicle-treated UCD males when compared to vehicle-treated non-UCD controls; and an almost significant increase in ATP [t=1.867, MD±SEM=-931.1±498.8, p=0.0775] in corticosterone-treated UCD animals when compared to corticosterone non-UCD controls (see Figure 3N-O).

### 3.6 Relationships between behavioural and biological phenotypes

Simple linear regression revealed several significant relationships between behaviour, inflammation, glucose metabolism, and weight change that survived correction for multiple comparisons. Distance travelled in the last OFT session before euthanasia were significantly negatively associated with weight change over five weeks [F(1,46)=24.78, R^2^=0.350, p<0.0001], IL1B gene expression in the NAc [F(1,44)=6.621, R^2^=0.131, p=0.0269], and TNFA gene expression in the NAc [F(1,43)=4.759, R^2^=0.100, p=0.0347]; and significantly positively associated with adipic acid concentration in the NAc [F(1,45)=8.185, R^2^=0.154, p=0.0190] (see Figure 4A-D). Rearing was significantly negatively associated with weight change [F(1,46)=18.20, R^2^=0.284, p=0.0003] and IL1B gene expression [F(1,44)=4.521, R^2^=0.093, p=0.0391], and significantly positively associated with adipic acid concentration in the NAc [F(1,45)=8.351, R^2^=0.157, p=0.0118] (see Figure 4E-G).

**Figure 4:**
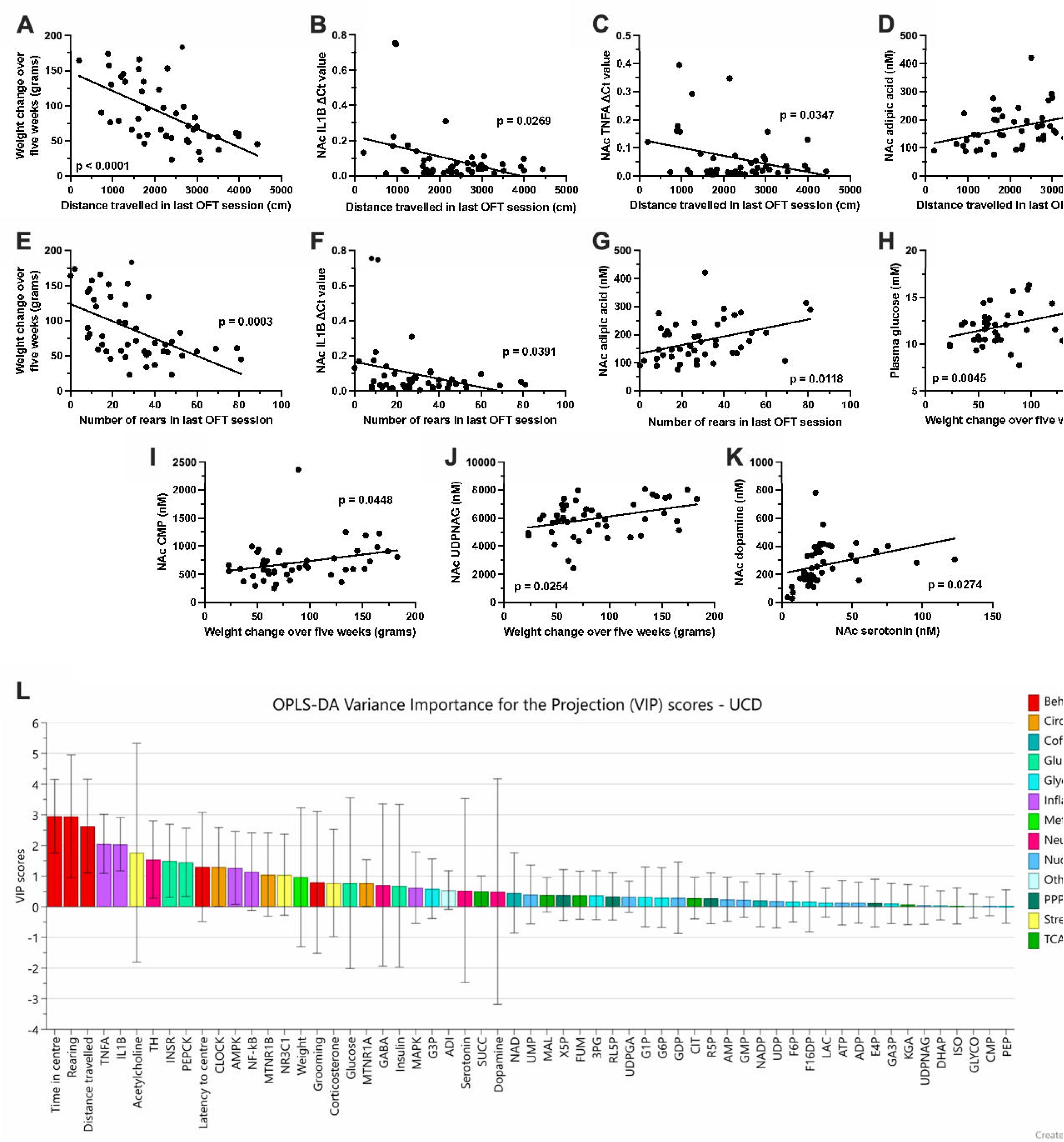
Correlations between behavioural and biological results. Scatterplots and linear regression models of distance travelled (cm) in the last open field test (OFT) session and weight change over five weeks (grams) ***(A)***, distance travelled and nucleus accumbens (NAc) IL1B gene expression (expressed as ΔCt) ***(B)***, distance travelled and NAc TNFA gene expression ***(C)***, distance travelled and NAc adipic acid concentration (nM) ***(D)***, number of rears in the last OFT session and weight change ***(E)***, number of rears and NAc IL1B gene expression ***(F)***, number of rears and NAc adipic acid concentration ***(G)***, weight change and plasma glucose (mM) ***(H)***, weight change and NAc cytidine monophosphate (CMP) concentration (nM) ***(I)***, weight change and NAc uridine diphosphate n-acetylglucosamine (UDPNAG) concentration (nM) ***(J)***, and NAc dopamine (nM) and NAc serotonin (nM) ***(K)***. Displayed p-values were calculated using simple linear regression followed by Holm-Šídák correction. Variance Importance in the Projection (VIP) scores of the orthogonal partial least squares discriminant analysis (OPLS-DA) model examining the effect of unpredictable circadian disturbance (UCD) ***(L)***. Variables are grouped and colour-coded according to marker type.

Weight change over five weeks was found to be significantly positively associated with plasma glucose [F(1,46)=11.39, R^2^=0.199, p=0.0045], NAc cytidine monophosphate [F(1,45)=4.259, R^2^=0.086, p=0.0448], and NAc uridine diphosphate n-acetylglucosamine [F(1, 45)=6.729, R^2^=0.130, p=0.0254] (see Figure 4H-J). Linear regression between neurotransmitter values also unveiled a significant positive relationship between dopamine and serotonin concentration in the NAc [F(1,45)=5.199, R^2^=0.104, p=0.0274] (see Figure 4K). These results suggest that UCD-induced inflammation, stress, disrupted glucose metabolism, and dysregulated circadian rhythm are highly interrelated.

### 3.7 Multivariate analyses

Multivariate analyses were conducted to identify whether the UCD, sex, and corticosterone groups were associated with specific patterns in biological/behavioural outcomes. According to R2X, R2Y, Q2 and ANOVA testing of cross-validated predictive residuals (CV-ANOVA) results, the model separated by sex had the strongest quality of fit, predictive validity, and model reliability (R2X=0.403; R2Y=0.852; Q2=0.565; CV-ANOVA: [F(1,47)=6.343, p<0.0001]), followed by UCD (R2X=0.116; R2Y=0.473; Q2=0.234; CV-ANOVA: [F(1,46)=6.719, p=0.0028]), and then corticosterone (R2X=0.114; R2Y=0.473; Q2=0.196; CV-ANOVA:[F(1,46)=5.34, p=0.0083]). Behavioural measures, inflammatory markers, stress markers, TH, glucose uptake markers, and circadian markers were found to contribute most to the UCD model (see Figure 4L). The aforementioned biological variables were more closely associated with UCD-treated animals, whereas the behavioural variables were more closely associated with non-UCD controls (see Supplementary Figure 3A). For the model based on sex, weight change, behaviour, corticosterone, neurotransmitters, and glucose uptake markers contributed most (see Supplementary Figure 3B). Weight change, latency to centre, glucose, GABA, and grooming were attributed to male animals, and distance travelled, corticosterone, rearing, time in centre, and corticosterone to females (see Supplementary Figure 3C). Finally, plasma corticosterone, glucose uptake markers, neurotransmitters, weight change, inflammatory markers, and CLOCK were found to be the highest contributing variables to the corticosterone model (see Supplementary Figure 4A). Neurotransmitter levels were more closely associated with the corticosterone group, and plasma corticosterone, glucose uptake markers, weight change, and inflammatory markers were more closely associated with vehicle-treated controls (see Supplementary Figure 4B).

## 4 Discussion

The role of circadian dysregulation in BD pathophysiology has been a focal point of research for decades (Takaesu, 2018). Several animal models of mania, such as the ClockΔ19 mutant (Roybal et al., 2007), paradoxical sleep deprivation (Gessa et al., 1995), and chronic unpredictable rhythm disturbance (Li et al., 2023), have attempted to capture elements of circadian dysfunction. However, no existing model has successfully replicated clinically-relevant circadian disturbances or induced mood-related behavioural shifts that mimic BD phenotypes (Beyer and Freund, 2017). In this study, we introduced the novel UCD protocol, designed to mirror key circadian disruptions observed in BD while minimising confounding effects on other physiological systems.

UCD was specifically developed to induce circadian fragmentation (through unpredictable light-dark cycling every two to three hours), poor sleep quality (via exposure to environmental noise), circadian misalignment (by varying the duration of light periods), and daytime dysfunction (using strobe lights to disrupt sleep during the dark phase or enforcing sleep through light exposure). These circadian disruptions are recognised as trait phenotypes of BD across all mood states. By effectively replicating these disturbances in a controlled setting, UCD may offer high translational value for studying psychiatric disorders characterised by dysregulation of arousal and general regulatory systems, including BD.

### 4.1 Behavioural effects

To assess mood-related behavioural alterations, we utilised locomotor activity metrics in the OFT. While locomotor behaviour can be influenced by multiple factors, it is a well-established, translational measure of mood dysregulation in preclinical models (Young et al., 2015). Manic episodes in BD patients are often characterised by increased exploratory activity and disorganised locomotor patterns, as observed in the human behavioural pattern monitor (Perry et al., 2009). Similarly, in rodents, rearing is considered a marker of exploratory drive (Perry et al., 2009), while reduced time in the centre of the arena (thigmotaxis) is associated with heightened anxiety and predator avoidance behaviours (Zhang et al., 2023). Conversely, increased time in the centre and higher centre-crossing frequency may indicate risk-taking behaviour, a hallmark of BD (Chen et al., 2010).

In the initial OFT session, when the environment was most novel, UCD-exposed animals exhibited increased exploratory and risk-taking behaviours compared to controls, suggesting heightened arousal and hyperactivity. These findings align with BD’s hyperarousal phenotype, supporting UCD’s relevance to manic-like behavioural states. Additionally, chronic corticosterone administration influenced behaviour, significantly increasing rearing frequency and total distance travelled, while having no effect on time in the centre or centre-crossing frequency. These effects are consistent with previous studies in which chronic corticosterone exposure (≥21 days) or chronic mild stress paradigms enhanced rearing behaviour and overall locomotor activity (Camargo et al., 2021; Harris et al., 1997; Spasojevic et al., 2016). Such findings further reinforce the hypothesis that stress system dysregulation contributes to BD-like arousal dysfunction.

To assess the temporal progression of behavioural changes, we employed repeated OFT. Habituation is an expected response in repeated behavioural testing, particularly in tasks reliant on novelty, such as the OFT (Chen et al., 2023). To account for this, we examined temporal behavioural stability across sessions. Non-UCD animals exhibited a U-shaped trajectory in rearing frequency and total distance travelled, a well-documented habituation pattern in rodent studies (Valle, 1971). In contrast, UCD-exposed animals displayed a distinct and negatively linear pattern, as identified through correlation analyses. Importantly, by the final week of testing, UCD animals exhibited a significant reduction in exploratory behaviour (rearing), indicative of hypoactivity. The consistency of this effect, combined with longitudinal analyses, suggests that these differences extend beyond simple habituation and reflect a fundamental shift in behavioural state. The significant change from a hypermotor and exploratory phenotype to a hypoactive state shown by UCD animals could reflect a “mood-shift,” which is a key phenotypic element of BD. This transition is particularly relevant given that mood-switching between manic and depressive states is a core feature of BD but is rarely recapitulated in preclinical models. The ability of UCD to induce this behavioural shift underscores its potential as a translationally relevant protocol for investigating BD pathophysiology.

There were clear differences in behaviour between males and females in this study. UCD-treated females, but not males, demonstrated persistent hyperactivity, with increased locomotor activity and exploratory behaviours that remained significant even in sex-stratified analyses. These findings align with previous research using the lateral hypothalamic kindling model of mania, which demonstrated that female rats displayed heightened rearing behaviour in the manic-like state compared to the post-manic phase, whereas males showed no significant differences between states (Abulseoud et al., 2015). Clinical studies indicate that female BD patients tend to exhibit more severe manic symptoms than their male counterparts, further supporting the translational relevance of these findings (Menculini et al., 2022). Conversely, UCD-treated males exhibited more pronounced habituation and an earlier transition to hypoactivity, with a reduction in locomotor behaviours occurring several days before UCD-treated females. A similar sex-specific effect has been observed in a rodent model of depression induced by myocardial infarction, where males and ovariectomised females developed depression-like behaviours, whereas intact females or those receiving estrogen replacement did not (Najjar et al., 2018). These findings suggest a potential neuroprotective role of ovarian hormones in modulating mood-related behaviour. However, in contrast to these preclinical findings, female BD patients report a higher lifetime burden of depressive symptoms and mixed episodes compared to males, highlighting the complexity of sex differences in BD pathophysiology (Morgan et al., 2005). These discrepancies may stem from hormonal, genetic, or environmental factors, underscoring the need for further research into sex-specific mechanisms of mood dysregulation.

### 4.2 Inflammatory, stress, and circadian markers

Inflammatory dysregulation is a well-established feature of BD, with increased levels of inflammatory markers observed across euthymic, manic, and depressive states (Poletti et al., 2024). Raised levels of IL-1β (Fiedorowicz et al., 2015; Pandey et al., 2015; Rao et al., 2010; Scaini et al., 2019; Soderlund et al., 2011), TNF-α (Bavaresco et al., 2020; Fiedorowicz et al., 2015; Kauer-Sant’Anna et al., 2009; Kim et al., 2007; Koga et al., 2019; Lee et al., 2021; Pandey et al., 2015; Uyanik et al., 2015), and NF-κB (Rao et al., 2010) have been observed in acutely ill and euthymic BD patients. While AMPK levels in BD patients have not been reported, its role in mitochondrial impairment suggests a potential contribution to BD pathophysiology (Morris et al., 2017). In this study, UCD-exposed animals exhibited increased expression of IL1B, TNFA, AMPK, and NFKB (in males only) in the NAc compared to non-UCD controls, suggesting an overall pro-inflammatory state in response to circadian disruption. However, since only gene expression was measured rather than protein levels, these results should be interpreted as indicative of a generalised inflammatory response rather than changes in specific cytokine activity.

Inflammation and locomotor activity appeared to be functionally linked, as NAc IL1B and TNFA expression were significantly negatively associated with rearing and total distance travelled in the final OFT session before euthanasia. These findings align with previous research demonstrating a direct correlation between inflammation and psychomotor retardation, both in healthy individuals exposed to inflammatory stimuli and in psychiatric patients (Goldsmith et al., 2016; Goldsmith et al., 2020). Strong associations have been reported between sleep disturbances and inflammation in BD, further supporting a mechanistic link between circadian dysfunction and immune activation (Dolsen et al., 2018; Fiedorowicz et al., 2015; Lee et al., 2021). Collectively, these results indicate that UCD-induced inflammation in the NAc mirrors key pathophysiological processes underlying BD.

Notably, inflammatory changes were observed only in vehicle-treated animals, suggesting that chronic corticosterone administration exerted an overall protective effect against inflammation. This is consistent with prior research showing that glucocorticoids exhibit both pro- and anti-inflammatory properties, depending on the context and duration of exposure (Smoak and Cidlowski, 2004). Interestingly, despite chronic corticosterone administration, plasma corticosterone levels were significantly lower in corticosterone-treated animals compared to vehicle-treated controls. This paradoxical effect has been reported in previous studies and is hypothesised to result from disrupted negative feedback regulation of the HPA axis following prolonged glucocorticoid exposure (Rosa et al., 2014). Although UCD did not significantly alter plasma corticosterone levels when sexes were analysed together, a sex-specific effect emerged upon stratification, with UCD-treated males showing a trend toward increased plasma corticosterone compared to non-UCD males. This suggests that UCD may enhance HPA axis activity, a phenomenon frequently observed in BD patients, particularly in those experiencing acute mood episodes (Deshauer et al., 2003; Fries et al., 2014; Mukherjee et al., 2022). The significant differences in biological and behavioural responses to UCD and chronic corticosterone administration, however, suggest that UCD’s effects cannot be attributed solely to HPA axis activation.

Further supporting the link between UCD, stress, and BD-like pathology, we observed significantly increased NAc acetylcholine levels in UCD-exposed animals. Acetylcholine has been shown to increase in response to stress (Mora et al., 2012). Notably, depressed BD patients exhibit heightened acetylcholine signalling, as evidenced by lower receptor availability across multiple brain regions compared to healthy controls (Hannestad et al., 2013). The convergence of acetylcholine dysregulation and increased inflammatory marker expression in UCD animals provides further support for a mechanistic link between circadian disruption, stress response, and mood dysregulation in BD.

Although melatonin levels in the NAc were undetectable using LC/MS-MS, UCD-treated animals displayed significantly increased expression of the melatonin receptor gene MTNR1B. This effect was restricted to corticosterone-treated animals; previous studies have demonstrated HPA axis-mediated regulation of melatonin signalling, supporting a potential interaction between stress and circadian pathways in BD pathophysiology (Gupta and Haldar, 2013). Additionally, UCD altered CLOCK gene expression, implicating core circadian transcriptional mechanisms in the observed behavioural and metabolic changes. The CLOCK/BMAL1 transcription factor complex has been shown to regulate melatonin receptor expression, further linking circadian and melatonergic signalling to UCD-induced disruptions (Beesley et al., 2015). These findings suggest that UCD affects circadian rhythm regulation at both transcriptional and receptor-mediated levels, reinforcing its translational relevance to BD.

The effects of UCD on inflammatory, stress, and circadian markers were markedly stronger in males than females. UCD-treated males exhibited increased plasma corticosterone, NAc acetylcholine, and upregulated expression of IL1B, TNFA, NFKB, CLOCK and MTNR1B gene expression compared to non-UCD male controls. In contrast, no significant differences in these markers were observed between UCD- and non-UCD females. Sex differences in neuroinflammatory responses have been reported in other animal models of mood disorders. For example, in a rodent model of depression induced by myocardial infarction, increased prefrontal cortex IL-1β, IL-2, IL-6, TNF-α, and IL-10 levels were found exclusively in depressed males, with no significant changes observed in females (Najjar et al., 2018). These findings suggest that males may be more susceptible to inflammation-driven mood dysregulation, which may partially explain the observed sex differences in UCD-induced inflammatory responses. Similarly, sex differences in HPA axis function have been well-documented in clinical studies. Depressed male patients exhibit higher serum and salivary cortisol levels, as well as greater cortisol reactivity, compared to depressed females, likely due to differences in receptor function and feedback sensitivity between sexes (Teo et al., 2023). Moreover, circadian rhythm regulation exhibits significant sex differences, including sex-specific patterns of clock gene expression, which may further influence mood regulation in BD (Bailey and Silver, 2014).

### 4.3 Metabolism

Circadian rhythms regulate multiple aspects of metabolism, including fat accumulation, insulin secretion, glucose homeostasis, and cellular respiration (Kim et al., 2019). Dysregulated metabolism is a hallmark of BD, and may be a major contributor to other important characteristics of the disease, such as insulin resistance, weight gain/loss, and changes in mood (Kim et al., 2019). Research has shown that obesity and type 2 diabetes occur at significantly higher rates in BD patients compared to the general population, further implicating metabolic dysfunction in the disorder’s pathophysiology (Miola et al., 2022).

In this study, UCD-treated animals were shown to have increased levels of NAc PEPCK and INSR gene expression when compared to non-UCD controls, which is considered to be indicative of insulin resistance (Qian et al., 2022). While plasma insulin and glucose levels were not significantly altered between UCD and non-UCD groups, this may have been influenced by the lack of fasting prior to euthanasia, which could have masked more subtle metabolic changes. Alternatively, the metabolic effects of UCD may be predominantly located within the brain and not extend to the periphery. Nonetheless, the observed alterations in NAc insulin signalling genes suggest that circadian disruption may impair metabolic regulation at the neural level, independent of systemic glucose changes.

UCD also had profound effects on body weight regulation. UCD-treated males gained significantly more weight than non-UCD controls, consistent with human studies showing that circadian disruption is a major risk factor for obesity (Noh, 2018). Furthermore, visceral adiposity has been implicated in both insulin resistance and systemic inflammation in BD patients, reinforcing the potential metabolic consequences of circadian misalignment (Miola et al., 2022). Interestingly, chronic corticosterone administration attenuated weight gain in male rats, an effect that has also been reported in rodent models of chronic mild stress (Harris et al., 1997). This suggests that chronic glucocorticoid exposure may exert protective effects against UCD-induced weight gain, possibly via alterations in energy expenditure, feeding behaviour, or metabolic rate. UCD-treated females displayed the opposite pattern to males, gaining significantly less weight than non-UCD female controls and failing to exhibit the same metabolic alterations as males. These findings do not align with clinical studies, which report higher rates of abdominal obesity in female BD patients compared to males (Baskaran et al., 2014). However, this discrepancy may be partially explained by sex differences in disordered eating, which is observed at higher rates in females with BD (Yakovleva et al., 2023).

The metabolomic profile of UCD-treated animals further supports a model of metabolic dysfunction. High-energy phosphates ATP and GDP were significantly reduced in the NAc of UCD-treated animals compared to non-UCD controls, suggesting a deficit in energy production within the limbic system. In contrast, corticosterone-treated animals exhibited increased levels of ATP and GDP, highlighting the divergent metabolic effects of stress hormone exposure versus circadian disruption. Further supporting this metabolic impairment, UCD-treated animals exhibited reductions in succinate and NAD, both of which play essential roles in oxidative phosphorylation. Succinate serves as a key intermediate in the citric acid cycle, directly contributing to ATP production (Tretter et al., 2016), while NAD functions as a critical cofactor in mitochondrial respiration and redox balance (Braidy et al., 2019). The observed reductions in these metabolites suggest that UCD-treated animals experience deficits in oxidative phosphorylation, a phenomenon that has also been documented in BD patients (Scaini et al., 2016). Regression analyses revealed significant negative associations between weight change and mood-related activity, reinforcing the interplay between metabolic function and arousal regulation. Collectively, these findings suggest that UCD-treated animals exhibit reduced oxidative phosphorylation, concurrent with changes in weight and hypoactivity, providing further evidence that circadian dysregulation directly affects energy metabolism and behavioural outcomes.

### 4.4 Limitations and future directions

This study provides key evidence that UCD can induce mood-related behavioural shift and concomitant biological changes relevant to psychiatric conditions involving general regulation and arousal systems dysfunction, such as BD. However, there are several limitations that need to be taken into consideration when interpreting experimental results. Firstly, sleep was not measured during this study. Sleep disturbance is an important phenotype of BD, and qualifying the effect of UCD on sleep is essential for fully understanding how the protocol affects circadian rhythm. It should be noted that environmental noise exposure, which was included in the UCD protocol, has been shown to increase sleep fragmentation and time spent awake in both female and male rats (Coborn et al., 2019). However, the other elements of UCD have unknown effects on sleep. Secondly, animals weren’t fasted before euthanasia, so the plasma glucose and insulin levels have limited interpretability. Also, food intake was not monitored, so it is unknown whether changes in appetite may have contributed to the weight and metabolic effects seen in UCD-treated animals. Thus, assessing the direct effects of UCD on sleep patterns and appetite would provide better knowledge about the mechanisms of the model. Furthermore, oestrous cycle measurements may aid in understanding the cause of sex-dependent differences induced by UCD. Investigating the effect of UCD on other dopaminergic and serotonergic markers within the NAc, such as those involved with uptake, release, or metabolism, may also be useful.

Further studies are needed to establish UCD as a translational animal model of dysregulated arousal and mood-shift. The repeated OFT was the only behavioural assessment conducted during this proof of principle study. It was chosen as a fast-screening tool to capture changes in mood-related behaviour, as temporal response and magnitude of mood-shift phenotype induced by UCD were unknown. Although changes in psychomotor activity are valid parameters for mood-related behaviours and mood-shift evaluation, further behavioural testing needs to be completed for fully characterise the face validity of UCD. Assessment of other behaviours (such as aggressiveness, sexual activity, and impulsivity) are necessary to consolidate the behavioural phenotyping. Biological samples should also be taken while animals are experiencing hyperactivity to determine any state-related differences in inflammation, metabolism, circadian rhythm, and stress. Finally, predictive validity needs to be assessed through the effectiveness of classical mood stabiliser medications, such as lithium, valproate, lamotrigine, and quetiapine.

## 5 Conclusion

This study provides comprehensive characterisation of the behavioural, inflammatory, circadian, stress, and metabolic effects of a novel UCD protocol, with and without chronic corticosterone administration. The findings highlight both overall and sex-dependent effects of UCD, supporting its translational relevance to BD and other psychiatric conditions involving dysregulated arousal and general regulatory systems.

Compared to non-UCD controls, UCD-treated animals exhibited a mood-related behavioural shift, initially displaying hyperactivity and manic-like behaviours, which then transitioned to a hypoactive state after four weeks of treatment. This pattern mirrors mood-state fluctuations in BD, reinforcing UCD’s potential as a preclinical model of mood dysregulation. At the molecular level, UCD animals showed significant increases in inflammatory markers (IL1B, TNFα), insulin resistance markers (INSR, PEPCK), circadian markers (MTNR1Β, CLOCK), and stress markers (acetylcholine, NR3C1) in the NAc. Additionally, UCD induced alterations in weight and central carbon metabolites, further linking circadian disruption to metabolic dysfunction.

Notably, sex-dependent differences were observed across both behavioural and biological variables. Females exhibited more pronounced behavioural differences between UCD and non-UCD groups, whereas males displayed greater biological alterations in response to UCD, particularly in inflammatory, circadian, and metabolic markers. These findings underscore the necessity of considering sex as a critical factor in preclinical studies of mood disorders. Furthermore, chronic corticosterone administration exerted a protective effect, mitigating UCD-induced inflammation, insulin resistance, and metabolic dysregulation. This suggests that glucocorticoid signalling may modulate the impact of circadian disturbances on stress and metabolic pathways, further emphasising the complex interplay between circadian disruption, stress response, and metabolic function in BD.

Together, these findings support the hypothesis that chronic circadian disturbance contributes to arousal dysfunction, a core pathophysiological feature of BD. By inducing mood-related behavioural shifts and neurobiological alterations relevant to BD, UCD represents a promising translational model for investigating circadian dysregulation in psychiatric disorders. Further validation through expanded behavioural assessments and pharmacological testing with mood stabilisers will be essential for establishing UCD’s utility as a preclinical model for BD pathophysiology.

## Acknowledgements

The authors would like to acknowledge the support and assistance provided by the University of Queensland Biological Resources Research Animal Facility (particularly Kym French), the Queensland Brain Institute Animal Behavioural Facility, and the University of Queensland Metabolomics and Proteomics (Q-Map) team for the completion of this study.

---

**Supplementary Table 1:**
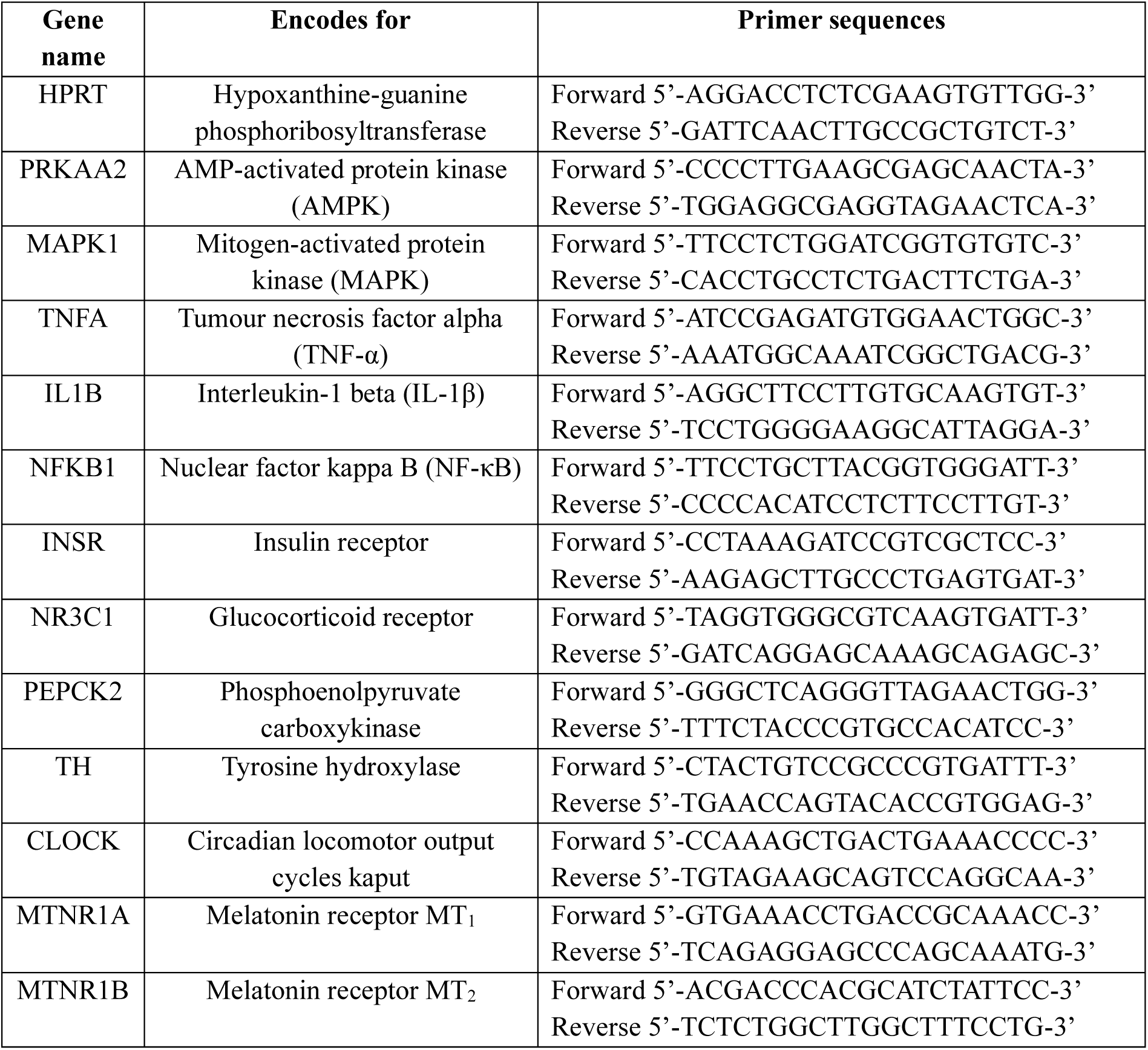
Primer details for real-time PCR.

**Supplementary Table 2:**
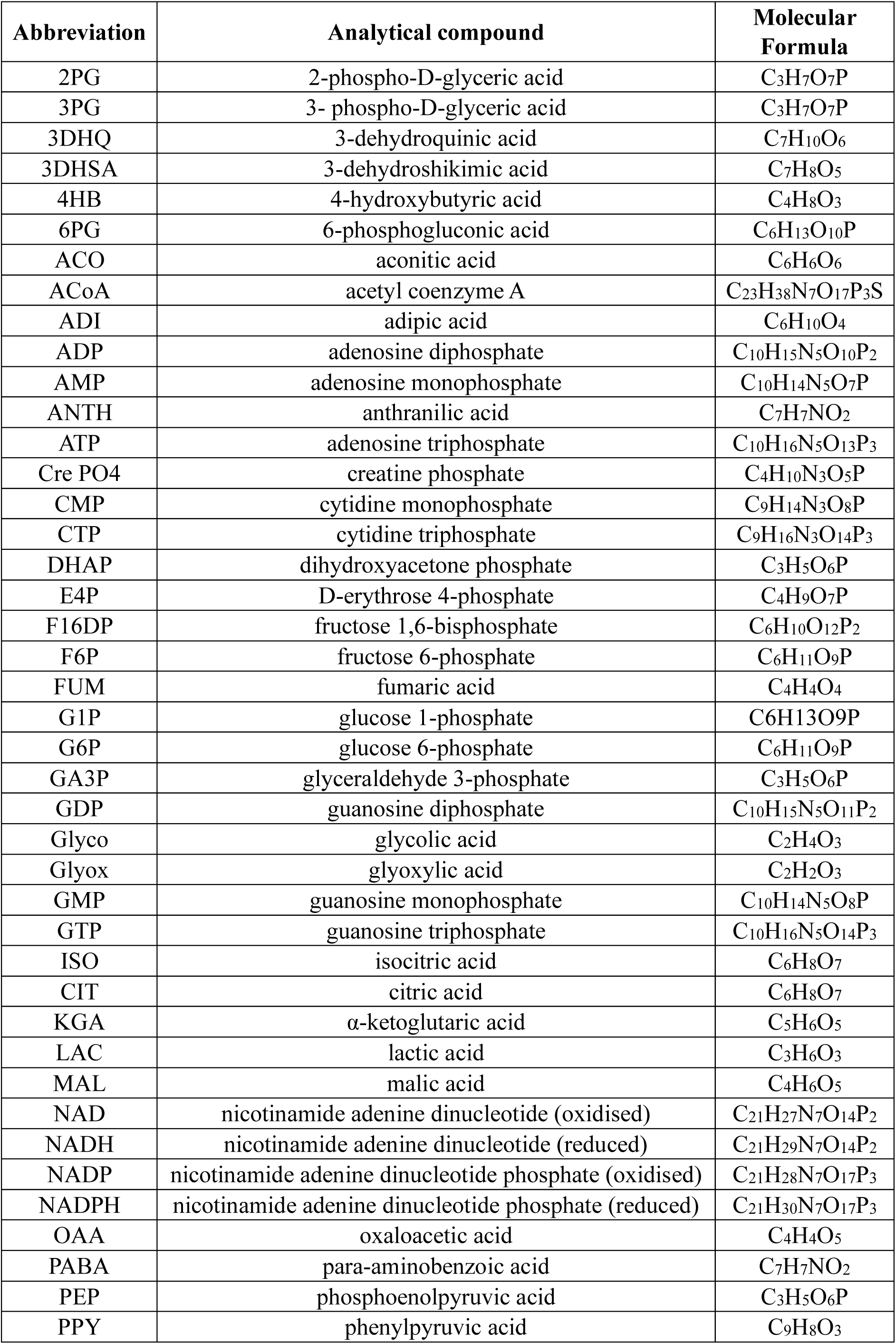
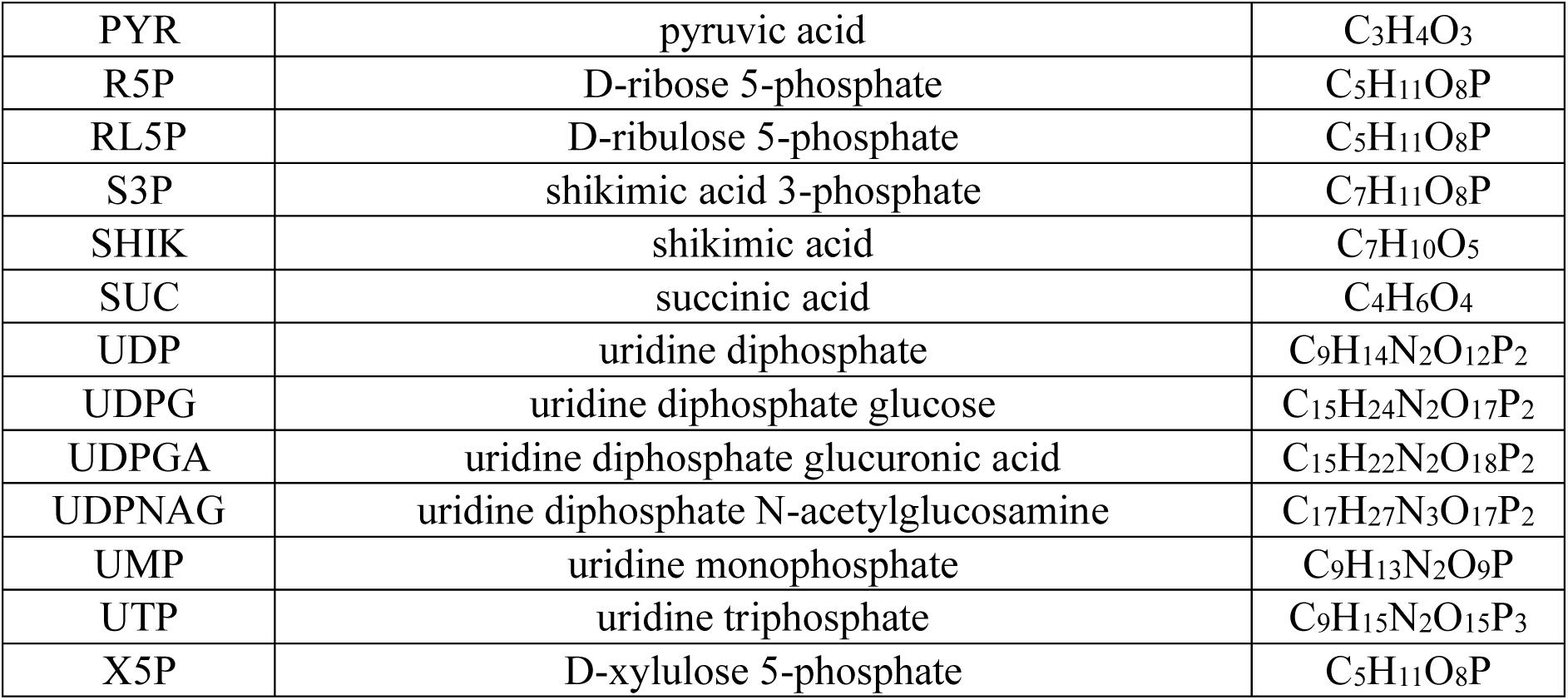
Central carbon metabolism (CCM) compound list.

| **Abbreviation** | **Analytical compound** | **Molecular Formula** |
| --- | --- | --- |
| 2PG | 2-phospho-D-glyceric acid | C<sub>3</sub>H<sub>7</sub>O<sub>7</sub>P |
| 3PG | 3- phospho-D-glyceric acid | C<sub>3</sub>H<sub>7</sub>O<sub>7</sub>P |
| 3DHQ | 3-dehydroquinic acid | C<sub>7</sub>H<sub>10</sub>O<sub>6</sub> |
| 3DHSA | 3-dehydroshikimic acid | C<sub>7</sub>H<sub>8</sub>O<sub>5</sub> |
| 4HB | 4-hydroxybutyric acid | C<sub>4</sub>H<sub>8</sub>O<sub>3</sub> |
| 6PG | 6-phosphogluconic acid | C<sub>6</sub>H<sub>13</sub>O<sub>10</sub>P |
| ACO | aconitic acid | C<sub>6</sub>H<sub>6</sub>O<sub>6</sub> |
| ACoA | acetyl coenzyme A | C<sub>23</sub>H<sub>38</sub>N<sub>7</sub>O<sub>17</sub>P<sub>3</sub>S |
| ADI | adipic acid | C<sub>6</sub>H<sub>10</sub>O<sub>4</sub> |
| ADP | adenosine diphosphate | C<sub>10</sub>H<sub>15</sub>N<sub>5</sub>O<sub>10</sub>P<sub>2</sub> |
| AMP | adenosine monophosphate | C<sub>10</sub>H<sub>14</sub>N<sub>5</sub>O<sub>7</sub>P |
| ANTH | anthranilic acid | C<sub>7</sub>H<sub>7</sub>NO<sub>2</sub> |
| ATP | adenosine triphosphate | C<sub>10</sub>H<sub>16</sub>N<sub>5</sub>O<sub>13</sub>P<sub>3</sub> |
| Cre PO4 | creatine phosphate | C<sub>4</sub>H<sub>10</sub>N<sub>3</sub>O<sub>5</sub>P |
| CMP | cytidine monophosphate | C<sub>9</sub>H<sub>14</sub>N<sub>3</sub>O<sub>8</sub>P |
| CTP | cytidine triphosphate | C<sub>9</sub>H<sub>16</sub>N<sub>3</sub>O<sub>14</sub>P<sub>3</sub> |
| DHAP | dihydroxyacetone phosphate | C<sub>3</sub>H<sub>5</sub>O<sub>6</sub>P |
| E4P | D-erythrose 4-phosphate | C<sub>4</sub>H<sub>9</sub>O<sub>7</sub>P |
| F16DP | fructose 1,6-bisphosphate | C<sub>6</sub>H<sub>10</sub>O<sub>12</sub>P<sub>2</sub> |
| F6P | fructose 6-phosphate | C<sub>6</sub>H<sub>11</sub>O<sub>9</sub>P |
| FUM | fumaric acid | C<sub>4</sub>H<sub>4</sub>O<sub>4</sub> |
| G1P | glucose 1-phosphate | C<sub>6</sub>H<sub>13</sub>O<sub>9</sub>P |
| G6P | glucose 6-phosphate | C<sub>6</sub>H<sub>11</sub>O<sub>9</sub>P |
| GA3P | glyceraldehyde 3-phosphate | C<sub>3</sub>H<sub>5</sub>O<sub>6</sub>P |
| GDP | guanosine diphosphate | C<sub>10</sub>H<sub>15</sub>N<sub>5</sub>O<sub>11</sub>P<sub>2</sub> |
| Glyco | glycolic acid | C<sub>2</sub>H<sub>4</sub>O<sub>3</sub> |
| Glyox | glyoxylic acid | C<sub>2</sub>H<sub>2</sub>O<sub>3</sub> |
| GMP | guanosine monophosphate | C<sub>10</sub>H<sub>14</sub>N<sub>5</sub>O<sub>8</sub>P |
| GTP | guanosine triphosphate | C<sub>10</sub>H<sub>16</sub>N<sub>5</sub>O<sub>14</sub>P<sub>3</sub> |
| ISO | isocitric acid | C<sub>6</sub>H<sub>8</sub>O<sub>7</sub> |
| CIT | citric acid | C<sub>6</sub>H<sub>8</sub>O<sub>7</sub> |
| KGA | α-ketoglutaric acid | C<sub>5</sub>H<sub>6</sub>O<sub>5</sub> |
| LAC | lactic acid | C<sub>3</sub>H<sub>6</sub>O<sub>3</sub> |
| MAL | malic acid | C<sub>4</sub>H<sub>6</sub>O<sub>5</sub> |
| NAD | nicotinamide adenine dinucleotide (oxidised) | C<sub>21</sub>H<sub>27</sub>N<sub>7</sub>O<sub>14</sub>P<sub>2</sub> |
| NADH | nicotinamide adenine dinucleotide (reduced) | C<sub>21</sub>H<sub>29</sub>N<sub>7</sub>O<sub>14</sub>P<sub>2</sub> |
| NADP | nicotinamide adenine dinucleotide phosphate (oxidised) | C<sub>21</sub>H<sub>28</sub>N<sub>7</sub>O<sub>17</sub>P<sub>3</sub> |
| NADPH | nicotinamide adenine dinucleotide phosphate (reduced) | C<sub>21</sub>H<sub>30</sub>N<sub>7</sub>O<sub>17</sub>P<sub>3</sub> |
| OAA | oxaloacetic acid | C<sub>4</sub>H<sub>4</sub>O<sub>5</sub> |
| PABA | para-aminobenzoic acid | C<sub>7</sub>H<sub>7</sub>NO<sub>2</sub> |
| PEP | phosphoenolpyruvic acid | C<sub>3</sub>H<sub>5</sub>O<sub>6</sub>P |
| PPY | phenylpyruvic acid | C<sub>9</sub>H<sub>8</sub>O<sub>3</sub> |
| PYR | pyruvic acid | $C_3H_4O_3$ |
| R5P | D-ribose 5-phosphate | $C_5H_{11}O_8P$ |
| RL5P | D-ribulose 5-phosphate | $C_5H_{11}O_8P$ |
| S3P | shikimic acid 3-phosphate | $C_7H_{11}O_8P$ |
| SHIK | shikimic acid | $C_7H_{10}O_5$ |
| SUC | succinic acid | $C_4H_6O_4$ |
| UDP | uridine diphosphate | $C_9H_{14}N_2O_{12}P_2$ |
| UDPG | uridine diphosphate glucose | $C_{15}H_{24}N_2O_{17}P_2$ |
| UDPGA | uridine diphosphate glucuronic acid | $C_{15}H_{22}N_2O_{18}P_2$ |
| UDPNAG | uridine diphosphate N-acetylglucosamine | $C_{17}H_{27}N_3O_{17}P_2$ |
| UMP | uridine monophosphate | $C_9H_{13}N_2O_9P$ |
| UTP | uridine triphosphate | $C_9H_{15}N_2O_{15}P_3$ |
| X5P | D-xylulose 5-phosphate | $C_5H_{11}O_8P$ |

**Supplementary Table 3:**
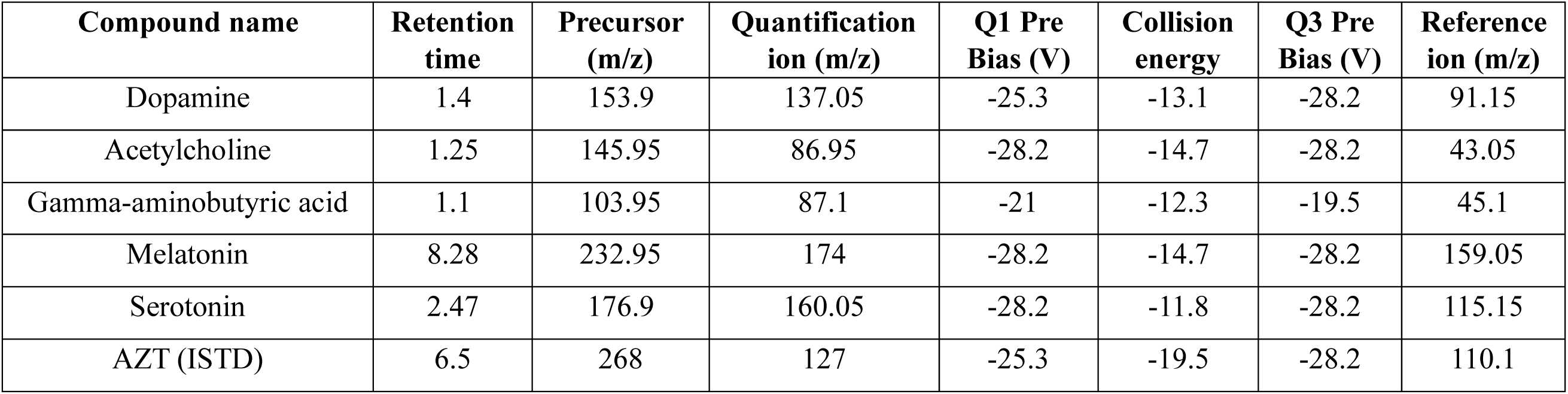
MRM transitions for quantifying neurotransmitters.

## Supplementary Data 2: *Statistical analyses*

### 2.10.1 Statistical analyses of behavioural data and weight gain

Three-way analysis of variance (ANOVA) with Geisser-Greenhouse correction was used on normal data to calculate differences between all groups in behavioural measures on each day of testing, and percentage difference from the first to final day of testing. Data were also separated into male-only and female-only; analysis of the same measures described above were performed on normal data using two-way ANOVA with Geisser-Greenhouse correction, and on non-normal data using multiple Mann-Whitney tests followed by the Holm-Šídák post-hoc test. Multiple comparisons were not performed on ANOVA analyses due to low power. Longitudinal analyses of behavioural data were conducted using Spearman’s correlation between behaviour on each day of testing and testing session (time), separated between groups.

### 2.10.2 Statistical analyses of plasma results, neurotransmitters, and gene expression data

Due to the predominantly non-normal nature of the data, three-way ANOVA analyses were not able to be performed. Instead, data from males and females were combined, and analyses on non-normal data were performed using multiple Mann-Whitney tests, and on normal data using multiple t-tests, both followed by the Holm-Šídák post-hoc test. Data were also separated into male-only and female-only and analysed in the same manner. Multiple comparisons were not performed on ANOVA analyses due to low power.

### 2.10.3 Analysis of metabolomics data

Three-way ANOVA with Geisser-Greenhouse correction was used on normal data to determine the effects of UCD or UCD x Corticosterone on metabolomics. Analyses were also performed using multiple t-tests on normal data or Mann-Whitney tests on non-normal data with males and females combined; and single t-test on normal data or single Mann-Whitney test on non-normal data with groups separated into UCD or non-UCD (n=24). Data were also separated into male-only and female-only; analysis of the same measures described above were performed on normal data using multiple t-tests or two-way ANOVA with Geisser-Greenhouse correction, and on non-normal data using multiple Mann-Whitney tests followed by the Holm-Šídák post-hoc test.

### 2.10.4 Analysis of relationships between behavioural and biological data

Simple linear regression analyses with Holm-Šídák correction for multiple comparisons were conducted to determine the relationships between behaviour in the final OFT session before euthanasia, weight, plasma measures, neurotransmitters, metabolomics, and gene expression. Statistical significance was set at p<0.05.

### 2.10.5 Multivariate analysis

Orthogonal partial-least squares discriminant analysis (OPLS-DA) was performed to identify which biological/behavioural variables differed most between UCD/non-UCD, male/female, and corticosterone/vehicle groups. OPLS-DA was chosen over PLS-DA as it is better suited to analyse noisy data. Variables were log-transformed and batch-scaled according to analysis type (behaviour, gene expression, plasma measures, neurotransmitter quantification, and CCM). Each model was cross-validated, and analysis of variance testing of cross-validated predictive residuals (CV-ANOVA) was performed to assess model reliability. All three models were found to be of acceptable quality of fit, predictive validity, and model reliability, as indicated by a R2X scores of less than 0.5, the difference between R2Y and Q2 being less than 0.3, and CV-ANOVA p-value scores of less than 0.05. OPLS-DA variable importance in the projection (VIP) scores were used to identify which variables drove separation between groups; variables with VIP scores over 1 were considered to be the highest contributors to the models.

---

**Supplementary Figure 1:**
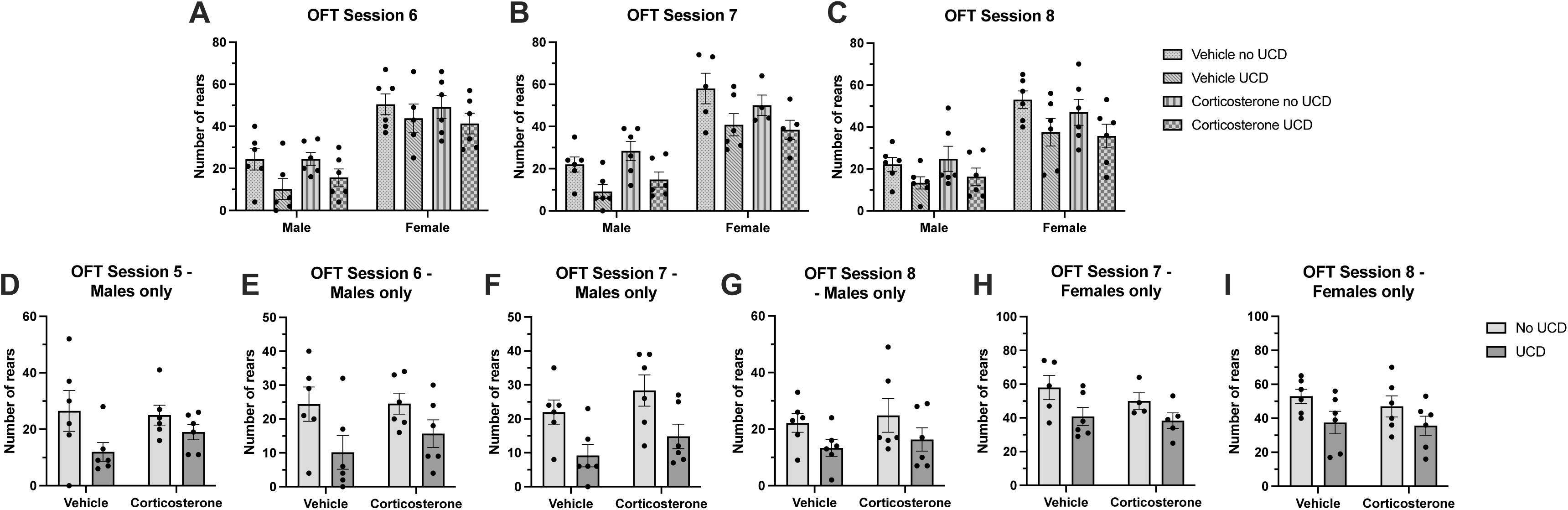
Effects of unpredictable circadian disruption (UCD) on locomotor behaviour. Mean (±SEM) number of rears on sessions six (**A**), seven (**B**), and eight (**C**) compared between UCD/non-UCD, corticosterone/vehicle, and male/female groups. Effect of UCD is p < 0.05 for analyses, as measured by three-way ANOVA with Geisser-Greenhouse correction. Mean (±SEM) number of rears on sessions five (**D**), six (**E**), seven (**F**), and eight (**G**) of males only; and number of rears on sessions seven (**H**) and eight (**I**) of females only, compared between UCD/non-UCD and corticosterone/vehicle groups. Effect of UCD is p < 0.05 for analyses, as measured by two-way ANOVA with Geisser-Greenhouse correction.

**Supplementary Figure 2:**
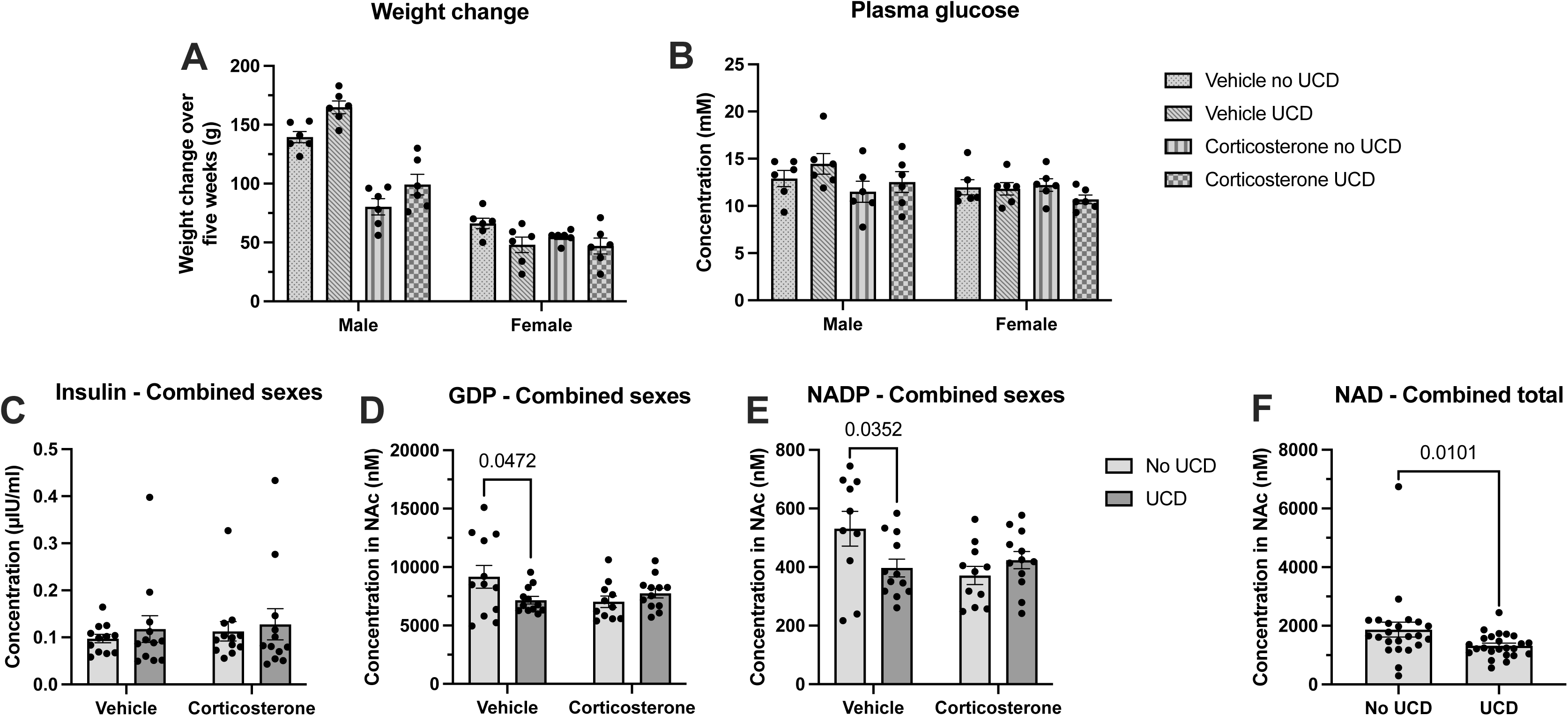
Effects of unpredictable circadian disruption (UCD) on weight gain and markers of glucose metabolism in plasma and the nucleus accumbens (NAc) Mean (±SEM) weight change over five weeks, compared between UCD/non-UCD and corticosterone/vehicle groups (**A**). Effect of corticosterone is p < 0.05, as measured by three-way ANOVA with Geisser-Greenhouse correction. Mean (±SEM) plasma glucose, compared between UCD/non-UCD and corticosterone/vehicle groups (**B**). Effect of Corticosterone and Gender x UCD is p < 0.10 as measured by three-way ANOVA with Geisser-Greenhouse correction. Mean (±SEM) plasma insulin (**C**), NAc guanosine diphosphate (GDP) (**D**), and NAc nicotinamide adenine dinucleotide phosphate (NADP) (**E**) of males and females combined, compared between UCD/non-UCD and corticosterone/vehicle groups. Mean (±SEM) NAc nicotinamide adenine dinucleotide (NAD) (**F**) of pooled UCD and non-UCD groups. Displayed p-values were calculated using multiple Mann-Whitney tests with Holm-Šídák correction.

**Supplementary Figure 3:**
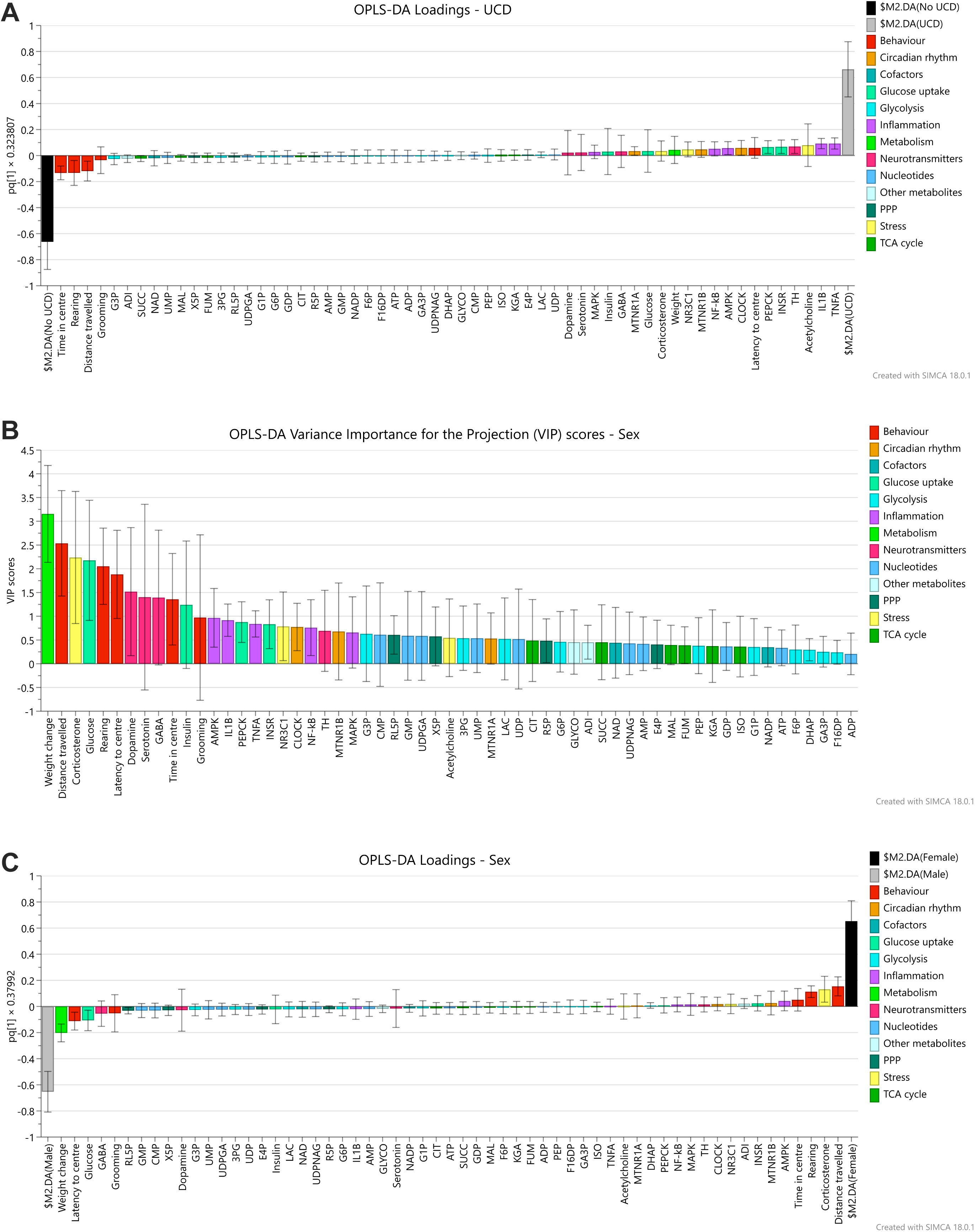
Orthogonal partial least squares discriminant analysis (OPLS-DA) results. Loading plot of the OPLS-DA model examining the effect of UCD (**A**). Variance Importance in the Projection (VIP) scores (**B**) and loading plots (**C**) of the OPLS-DA model examining the effect of sex. Variables are grouped and colour-coded according to marker type.

**Supplementary Figure 4:**
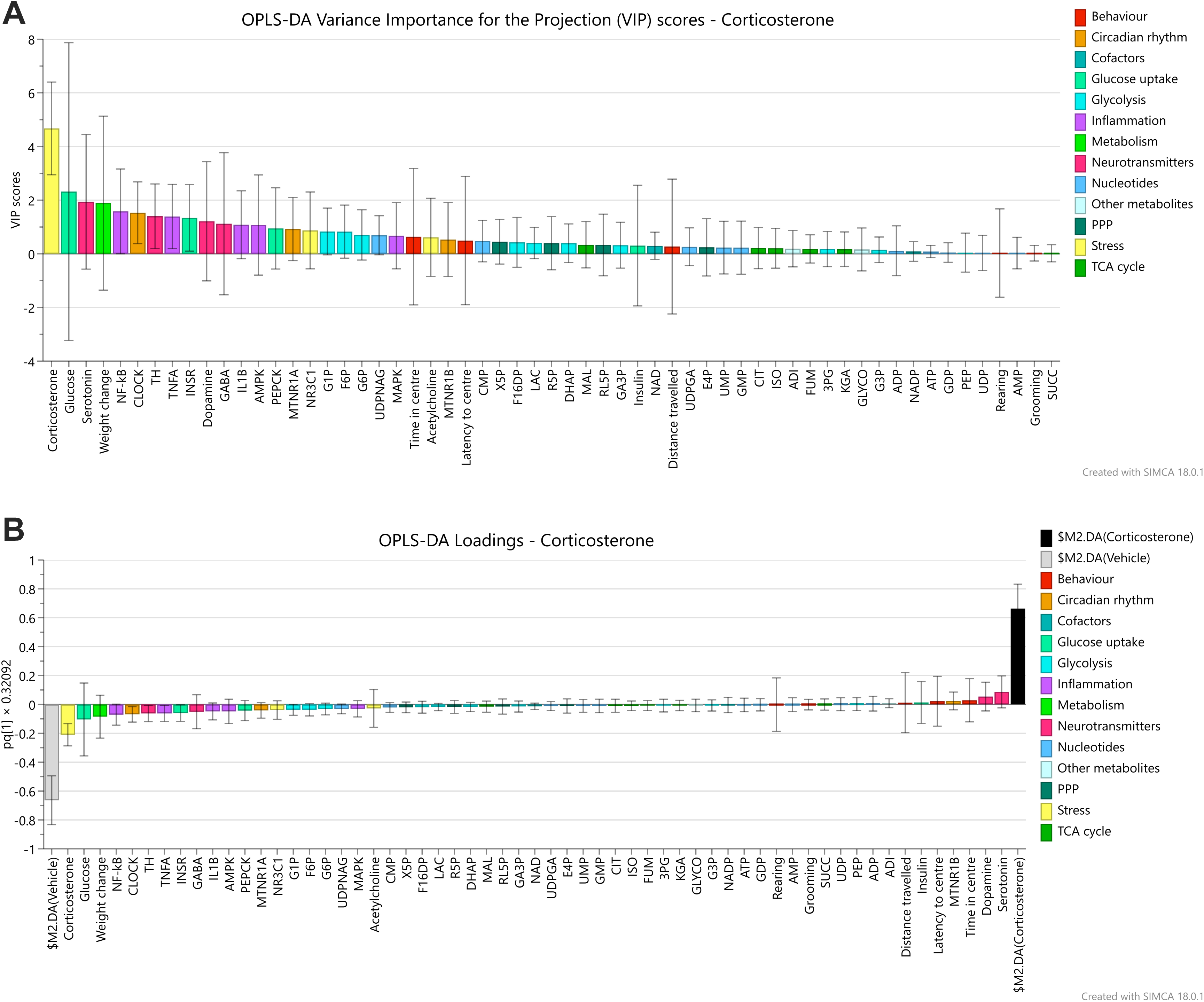
Orthogonal partial least squares discriminant analysis (OPLS-DA) results. Variance Importance in the Projection (VIP) scores (**A**) and loading plots (**B**) of the OPLS-DA model examining the effect of corticosterone. Variables are grouped and colour-coded according to marker type.

